# *Toxoplasma* Membrane Inositol Phospholipid Binding Protein TgREMIND Is Essential for Secretory Organelle Function and Host Infection

**DOI:** 10.1101/2023.03.19.533351

**Authors:** Rodrigue Houngue, Lamba Omar Sangare, Tchilabalo Dilezitoko Alayi, Aissatou Dieng, Tristan Bitard-Feildel, Claire Boulogne, Christian Slomianny, Cynthia Menonve Atindehou, Lucie Ayi Fanou, Yetrib Hathout, Jeroen PJ Saeij, Isabelle Callebaut, Stanislas Tomavo

## Abstract

Apicomplexan parasites have specialized secretory organelles called rhoptries, micronemes, and dense granules that are essential for host infection. Here, we show that TgREMIND, a *Toxoplasma gondii* protein containing a membrane phospholipid interacting domain, is required for the biogenesis of rhoptries and dense granules. TgREMIND contains a Fes/CIP4 homology-Bin/Amphiphysin/Rvs (F-BAR) domain at the N-terminus, known to promote cell membrane bending, and a novel uncharacterized domain that we named REMIND for <u>re</u>gulator of <u>m</u>embrane <u>in</u>teracting <u>d</u>omain at the C-terminus. TgREMIND binds to PIP2 lipid species and both F-BAR and REMIND domains are necessary to ensure proper biological activities *in vitro* and *in cellulo*. Conditional depletion of TgREMIND results in the absence of dense granules and abnormal transparent rhoptries, leading to a severe inhibition of parasite motility, host invasion, and dissemination. Thus, our study demonstrates that TgREMIND is essential for the proper functioning of key secretory organelles required for successful infection by *Toxoplasma*.

## INTRODUCTION

*Toxoplasma gondii* (*T. gondii*) is an obligate intracellular eukaryotic parasite of the phylum *Apicomplexa*, which also includes the deadly human malaria pathogen *Plasmodium falciparum*. Toxoplasmosis is principally dangerous to fetuses of primo-infected pregnant women and individuals with compromised immune systems. Infection by *T. gondii* is life-long and incurable, leaving chronically infected people susceptible to reactivated disease, including severe ocular disorders and vision loss. A better understanding of the cellular and molecular mechanisms of infection is key to uncovering new therapeutic strategies against *T. gondii*. After entry inside the host cell, *T. gondii* establishes its permissive replication niche, named the parasitophorous vacuole (PV), which is demarcated from the host cytoplasm, by a membrane (PVM) (Wang et al., 2020). To develop inside the host cell, apicomplexan parasites possess unique morphological features that constitute the hallmark of the phylum (Gubbels and Duraisingh, 2012; Tagoe et al., 2021). Among these features, is the remarkable apical complex composed of the polar ring, the conoid and lineage-specific secretory organelles called rhoptries and micronemes that play an essential role in apicomplexan parasite pathogenesis. In addition, a third secretory organelle called dense granules releases GRA proteins that are involved in the formation of the PV, in which the parasites replicate intracellularly, and in several other functions, such as transfer of small molecules from the host cytoplasm into the PV (Gold et al., 2015; Wang et al., 2020). The molecular mechanisms and functions played by these organelles during host cell attachment, invasion and survival are described in more detail elsewhere (Huynh et al., 2003; Harper et al., 2006; Saeij JP et al., 2007; Boothroyd and Dubremetz, 2008; Butcher et al., 2011; Griffith et al., 2022). However, the underlying mechanisms involved in the biogenesis of these organelles and their maintenance throughout the parasite replication cycle remain incompletely understood. The secretory organelles are synthesized *de novo* during the replication and budding of the daughter parasites formed within the mother (Tomavo et al., 2013). Moreover, transport of ROP and MIC proteins depends on their appropriate timing of expression, which suggests that their intracellular trafficking is directly associated with secretory organelle biogenesis (Ngo et al., 2003, Harper et al., 2006). However, more information is needed about how the parasite-specific factors that supply these secretory organelles are differentially sorted and transported in a timely, coordinated, and regulated fashion. We described that forming these vital secretory organelles requires *T. gondii* sortilin-like receptor TgSORTLR, also named TgSORT (Sloves et al. 2012; Honfozo et al., 2023). The N-terminal and luminal ectodomain of the receptor binds ROP and MIC proteins while the cytosolic tail binds to the TgVps35-retromer complex to recruit partners to enable anterograde and retrograde receptor transport (Sangaré et al. 2016). Here, we describe a new parasite-specific protein TgREMIND, which can drive membrane bending, deformation, and vesicle budding through interactions with inositol phospholipids. TgREMIND contains a highly conserved Fer-CIP4 homology-BAR (F-BAR) domain at the N-terminus and a novel uncharacterized REMIND domain (for <u>re</u>gulator of <u>m</u>embrane <u>in</u>teracting <u>d</u>omain) at the C-terminus. The presence of both F-BAR and REMIND domains in this protein, which is therefore named TgREMIND, are necessary to ensure efficient biological activities *in vitro* and *in cellulo*.

Conditional targeted ablation demonstrates that mutants devoid of TgREMIND lose dense granules and typical opaque rhoptries, leading to severe inhibition of host cell invasion. Our findings illustrate the key role of TgREMIND, a lipid-binding protein, in the formation and functions of secretory organelles required for host infection and parasite dissemination.

## RESULTS

### Identification of Molecular Features Associated with Functions of TgREMIND

We first used computational analysis to identify potential parasite proteins that could interact with lipids, deform membranes and create vesicles containing proteins/lipids destined to the parasite’s secretory organelles. This analysis identified TgREMIND, a new *T. gondii* regulator containing a membrane-interacting domain, as a potential candidate. TgREMIND was one of forty hypothetical proteins co-immunoprecipitated with TgVps35-cMyc using a total protein extract from our TgVps35-cMyc strain, which was made in the background of the conditional iKO-TgSORT mutant. TgREMIND is composed of two foldable domains, separated by a large disordered region (Figure S1A). The first foldable domain (aa 79-369) shares significant similarities with the F-BAR (FES-CIP4 Homology and Bin/Amphiphysin/Rvs) domains of known 3D structures, such as the tyrosine-protein kinase Fes/Fps (pdb 4DYL), formin-binding protein 1 (pdb 2EFL), and septation protein Imp2 (pdb 5C1F) F-BAR domains. We found this F-BAR domain in several homologs of TgREMIND in different apicomplexan parasites (Figure S2). The second foldable domain (aa 507-825), which we named REMIND, does not share obvious similarities with sequences of known 3D structures, but its middle part (aa 588-749) matches the Pfam SET binding factor 2 (SBF2) profile. A search of the UniProt reference proteomes using PSI-BLAST identified significant similarities between the sequence of the C-terminal region of REMIND and proteins of *Alveolata* and Oomycetes species, where it is also associated with an N-terminal F-BAR domain (Figures 1A and S3). The REMIND C-terminal region was also detected in other eukaryotic proteins, associated with different domain architectures, such as TBC (Tre-2, BUB2p, and Cdc16p) domains characteristic of Rab GAPs. It is also linked to the upstream u-DENN (differentially expressed in normal and neoplastic cells), core c-DENN, and downstream d-DENN domains (characteristic of Rab GEFs) in human MADD as well as in human MTMR5 and MTMR13, both members of the myotubularin-related protein family (Figures 1A and S3).

**Figure 1.**
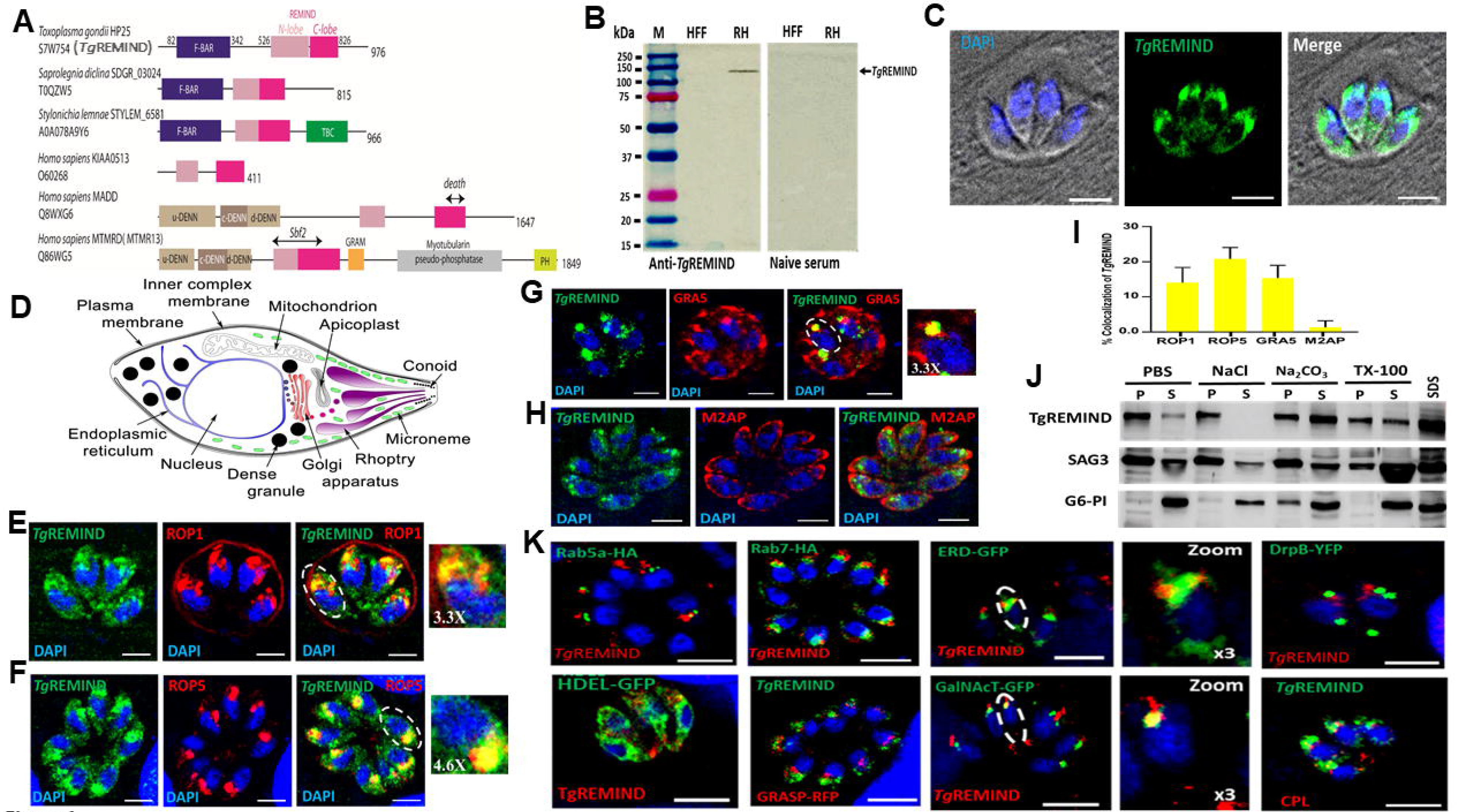
TgREMIND is a membrane-bound F-BAR-containing domain protein present in different subcellular compartments. A) Domain architecture of representative members of the REMIND family. B) Immunoblots probed with mouse polyclonal anti-TgREMIND antibodies using *T. gondii* (RH strain). Human foreskin fibroblasts (HFF) and naive sera showed no protein band, as expected. C) Localization of TgREMIND using anti-TgREMIND antibodies and confocal microscopy. D) Schematic representation of *T. gondii*. E and F) TgREMIND partially co-localizes with ROP1 and ROP5 proteins; G) TgREMIND also partially co-localizes with GRA5 protein. H) TgREMIND shows no co-localization with microneme M2AP. I) Quantification of TgREMIND co-distribution with ROP1, ROP5, GRA5 and M2AP. Bars represent n=3 independent experiments ± standard deviation (SD). J) Cell fractionation on wild type *T. gondii* parasites and Western blots using TgREMIND, GPI-anchored surface Antigen 3 and the glycolytic enzyme, Glucose 6-Phosphate Isomerase (G6-PI). K) Confocal images of intracellular parasites transiently transfected with plasmids expressing markers of different compartments and organelles and probed with anti-TgREMIND antibodies. Bars, 3 µm.

### TgREMIND Resides in the Cytoplasm and close to Rhoptries and Dense Granules

We produced specific mouse polyclonal antibodies against recombinant GST-tagged full-length TgREMIND (aa_1_-_976_), the F-BAR (aa_1-386_), and REMIND (aa_387_-_976_) domain. These polyclonal antibodies recognized a single band of 140 kDa (Figure S4A), which was exclusively expressed in parasites and not in uninfected human fibroblasts (HFF) (Figure 1B). However, this protein size is larger than the predicted molecular mass of TgREMIND (TGGT1_259720, see www.toxodb.org), estimated at 100 kDa. We also appended an encoded c-Myc tag to TgREMIND using a knock-in strategy, which ensures steady-state levels of epitope-tagged protein expression via homologous promoters (Huynh and Carruthers, 2009). Again, the anti-cMyc antibodies (Figure S4B) recognized a band of 140 kDa. This difference in molecular size could be due to the chemical nature and behavior of the amino acids of TgREMIND or the presence of a stretch of several 3 and 5 serine residues at the C-terminus of the protein (Figure S1B), suggesting multiple possible post-translational modifications. Confocal imaging using anti-TgREMIND (Figure 1C) and anti-cMyc (Figure S4C, top left panel) antibodies revealed labeling of the cytoplasm of *T. gondii* with a stronger signal at the apical region above the nucleus. This fluorescence appeared close to TgSORTLR, TgVps35, and proROP4 (Figure S4C). Based on the location of different compartments and organelles of *T. gondii* drawn by the schematic representation in Figure 1D, TgREMIND partially co-distributed with ROP1 and ROP5 (Figure 1E and 1F), with the strongest co-distribution observed with ROP7 (Figure S4D). Quantifying these confocal images showed that 15 ± 3% of ROP1 and 20 ± 5% of ROP5 protein co-distributed, respectively, with TgREMIND (Figure 1I). In addition, we found that 17 ± 3% of GRA5 co-distributed with TgREMIND (Figure 1G and 1I). In contrast, there was no co-distribution between the microneme M2AP protein and TgREMIND (Figure 1H and 1I). Together with the presence of the F-BAR domain, these data suggest a close vicinity of TgREMIND to different intracellular membrane compartments in *T. gondii*.

### TgREMIND Interacts with Cellular Membrane Components and Secretory Organelles

To corroborate the location of TgREMIND with its biochemical properties, we lysed extracellular parasites with PBS alone or PBS containing either detergent or other chaotropic agents, and separated insoluble proteins (P) from the soluble (S) fractions. Under these conditions, TgREMIND was insoluble in PBS alone and in PBS containing 0.5M NaCl but partially soluble by detergent extraction (2% TritonX-100) and much more extracted by sodium carbonate (Na_2_CO_3_) (Figure 1J). In contrast, detergent (Triton X-100) extracted the positive control membrane GPI-anchored Surface Antigen 3 (GPI-SAG3), whereas the cytosolic glycolytic enzyme glucose-6 phosphate isomerase (G6-PI) was soluble in PBS, as expected (Figure 1J). As sodium carbonate extraction is a canonical way to distinguish integral membrane proteins from other hydrophobic moderately membrane‐associated proteins (Kim et al. 2015), these data indicate that a significant amount of TgREMIND binds to subcellular membranes that are present in the cytoplasm of *T. gondii*. Furthermore, using other subcellular markers of *T. gondii*, we observed weak co-localization with post-Golgi (ERD-GFP and N-acetylgalactosaminyltransferase (GalNAc-T)), and late endosomal-like compartment such as Rab7-HA (Figure 1K), indicating that TgREMIND is present in distinct subcellular compartments of the parasite simultaneously. However, no co-localization was found with the ER (HDEL-GFP), Golgi (GRASP-GFP), and dynamin-related compartment (DrpB-GFP), suggesting that TgREMIND does not interact with these later compartments (Figure 1K).

To gain further insights into the nature and location of cellular membrane proteins interacting with TgREMIND, we pulled down its partners using purified recombinant GST-full length (FL) TgREMIND, its F-BAR or REMIND domains, and total parasite protein extracted by Triton-X 100, followed by mass spectrometry analyses. As shown in Figure S5A, all three or two baits share 118 proteins (52%) of the 224 proteins pulled down. However, GST-REMIND, GST-(FL) TgREMIND, and GST-F-BAR alone pulled down only 19 (8%), 28 (12%), and 59 (26%) proteins, respectively. Interestingly, we identified numerous proteins, which were categorized as follows in Table S1: A) Fifty-five factors involved in lipid/protein trafficking and biogenesis of vesicles and organelles; B) Twelve rhoptry (ROP) proteins; C) Twelve dense granule (GRA) proteins; D) Four microneme (MIC) proteins; and E) Twenty-six cytoskeleton and microtubule components. This functional classification of all proteins identified by mass spectrometry revealed that some of the interactors played roles in membrane traffic of vesicle membrane from ER to Golgi apparatus, recycling from ELC to post-Golgi, organelle biogenesis, and parasite motility in addition to regulators of membrane lipids, transporter/exchangers of membrane lipids and other trafficking vesicle formation (Table S1). As several proteins of rhoptries and dense granules were identified, these data agree with the co-distribution observed between TgREMIND with ROP and GRA proteins, while no co-distribution was noticed with microneme MIC proteins. We conclude that TgREMIND, with its membrane lipid interacting and regulator F-BAR and REMIND domains, may be involved in the formation of rhoptries and dense granules.

### TgREMIND Silencing Abrogates Secretion of ROP and GRA Proteins

We conditionally knocked out TgREMIND gene using the strategy in Figure 2A. We selected two positive knockout clones, which showed a perfect deletion of the endogenous target gene (Figure 2B). Treatment with anhydrotetracycline (ATc) resulted in the disappearance of the TgREMIND protein in the two knockout clones (iKO1 and iKO2), as revealed by Western blots (Figures 2C and 2D) and confocal microscopy (Figure 2E). We complemented this mutant with the full-length FLAG-tagged-TgREMIND (Figure 2F and 2G, see Comp-iKO2), which was introduced in the non-essential uracil phosphoryl transferase (UPRT) locus (Fox and Bzik, 2002). We confirmed the complementation of iKO-TgREMIND mutants as HA-TgREMIND, and TgREMIND-FLAG proteins were simultaneously expressed (Figure 2 Merge F+G, see the yellow band). Only TgREMIND-FLAG protein was present under ATc treatment (Figure 2 Merge F+G, see the red band). Under ATc pressure, TgREMIND deficiency revealed the complete absence of five distinct dense granule proteins (GRA1, GRA2, GRA3, GRA4, and GRA5) in the PVM and intra-vacuolar space of the intracellular mutants (Figure 3A, right images) as compared to decoration of the PVM by these GRA proteins when these mutants were grown in the absence of ATc (Figure 3A, left images). Furthermore, the ROP1 protein was absent in the PVM of these mutants (Figure 3B, top right and middle images). As expected, untreated parasites displayed abundant staining of the PVM by ROP1 (Figure 3B, top left and middle images). The signal of ROP7 protein, which is not normally present in the PVM, was unchanged regardless of ATc treatment (Figure 3B, bottom left and right panels). Complementation of the iKO-TgREMIND mutants fully restored the secretion of both GRA and ROP proteins into the PVM in the presence of ATc (Figure 3C). Biochemical analyses showed that depletion of TgREMIND after ATc pressure led to a ∼70% reduction of secretion of GRA1 protein from extracellular mutants that were incubated with propranolol (Bullen et al., 2016), known to trigger the release of GRA and MIC proteins in the extracellular medium (Figures 4A and 4B). In contrast, no difference was observed in microneme MIC9 protein regardless of ATc and propranolol treatment (Figures 4A and 4C). Furthermore, we confirmed that the apical fluorescence pattern of three other micronemes, M2AP, MIC2, and MIC5, was unchanged in ATc-depleted or untreated TgREMIND mutants (Figure 4D, right and left panels). Confocal imaging showed that all other intracellular organelles, such as the apicoplast (API), the mitochondrion (HSP60), CPL-containing organelle (known to be involved in the maturation of MIC proteins), the inner membrane complex (IMC), the centriole (CEN) and plasma membrane (SAG1) were not affected by the depletion of TgREMIND protein in these mutants (Figure 4E). These data indicate that the suppressive effect of TgREMIND deletion on protein secretion into the PVM is restricted and specific to dense granules and rhoptries.

**Figure 2.**
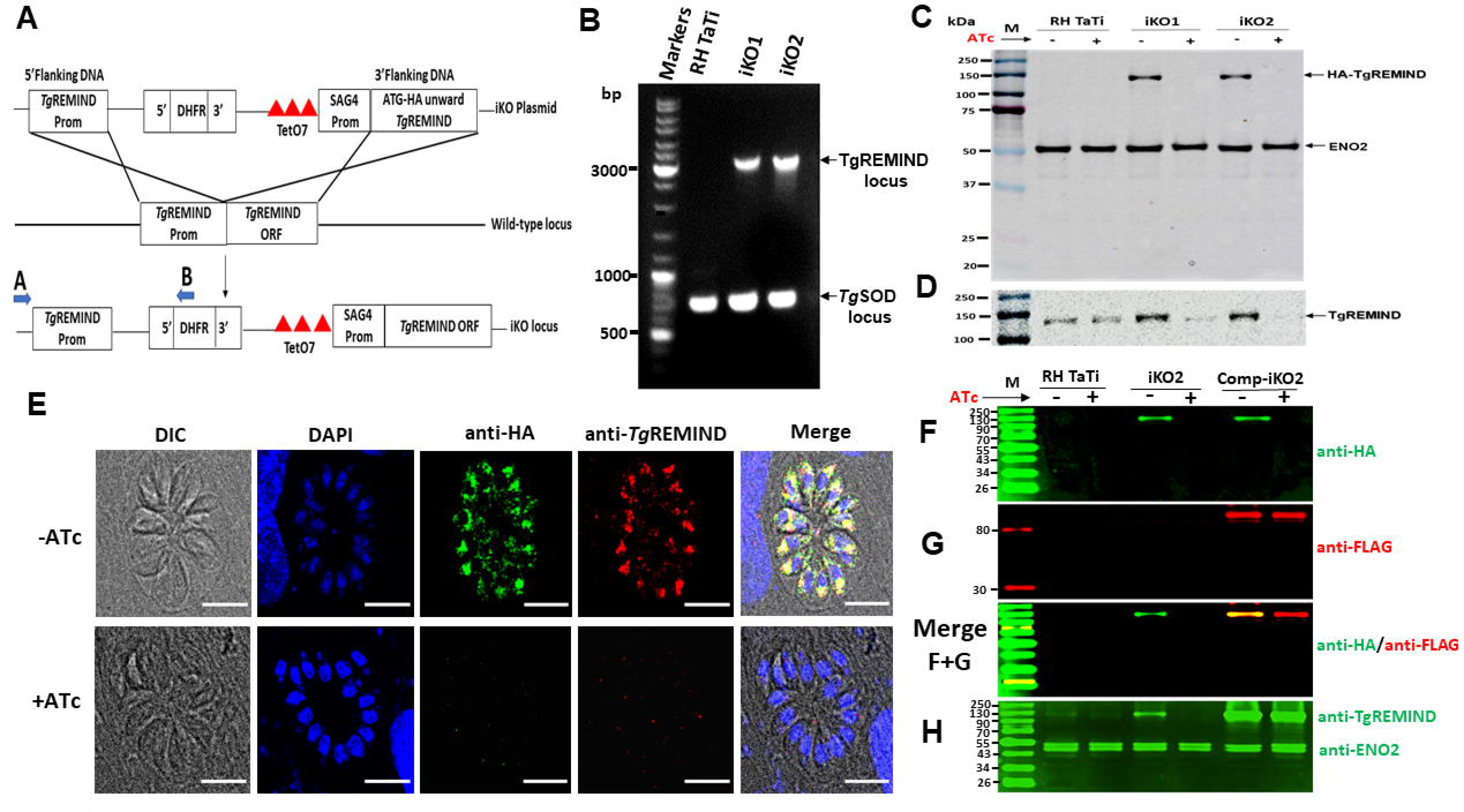
Conditional ablation of TgREMIND gene and complementation. A) Schematic of the approach used for the conditional ablation of TgREMIND gene. B) PCR analysis using the primers A and B (see blue arrows in panel A) confirms the conditional targeted ablation of TgREMIND. *T. gondii* superoxide dismutase (TgSOD), positive control. C) Immunoblots of wild type parasites and two clones (iKO1 and iKO2) of TgREMIND-deficient lines grown in the presence or absence of ATc for 48 h and probed with anti-HA antibodies. ENO2 protein, a loading control. D) Immunoblots, as in panel C, but probed with anti-TgREMIND antibodies. E) Confirmation of TgREMIND conditional depletion by confocal imaging using anti-HA and anti-TgREMIND antibodies. Bar, 3 µm. F) Immunoblots of untreated or treated wild type, iKO-TgREMIND and complemented Comp-iKO2 probed with anti-HA antibodies and secondary antibodies Alexa Fluor 647 nm goat anti-rat. G) Immunoblots probed with anti-FLAG antibodies and secondary antibody Alexa Fluor plus 800 nm goat anti-rabbit. F+G) Merged of panels F and G. H) Immunoblots with the loading materials as in panel A but probed with anti-TgREMIND and ENO2 (loading control) antibodies.

**Figure 3.**
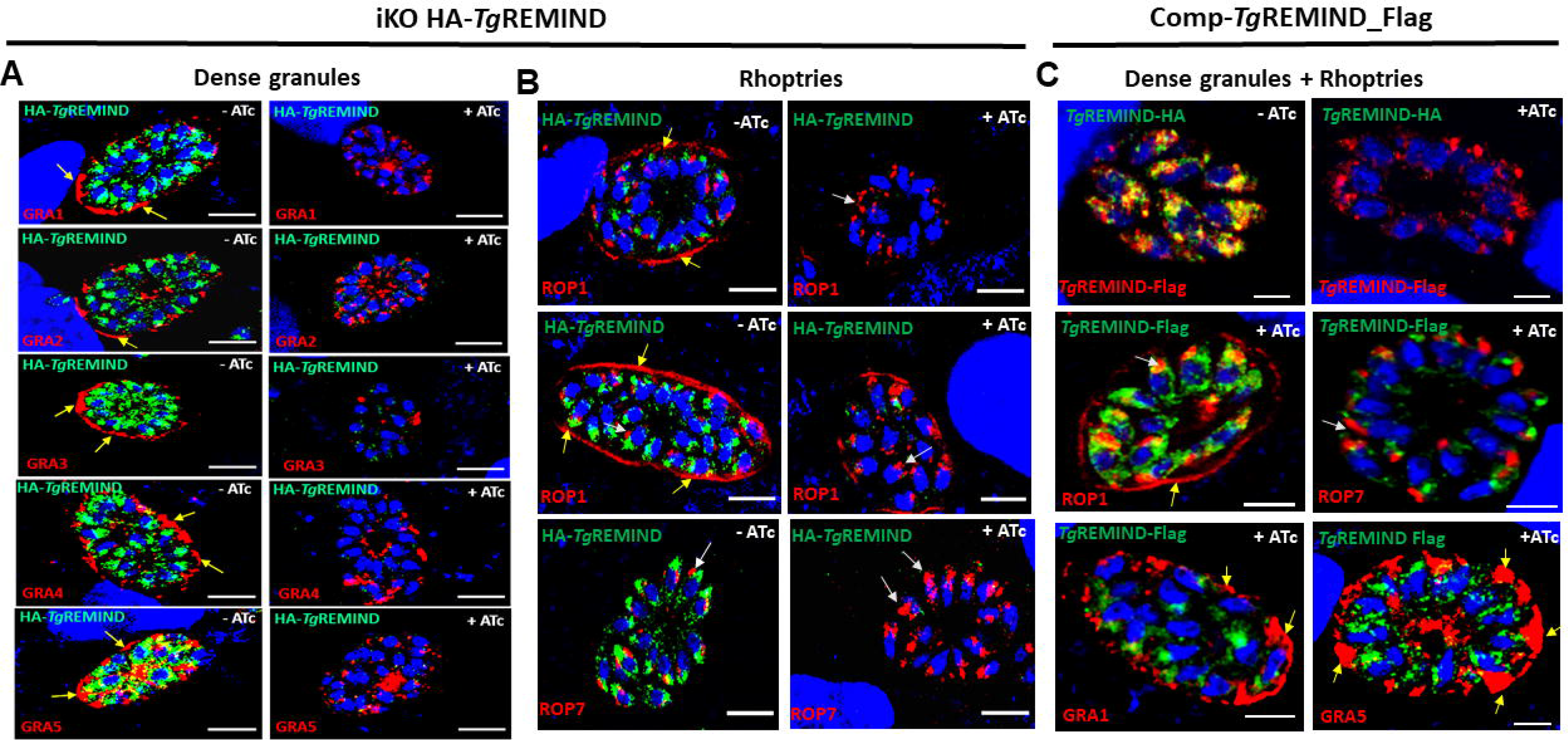
The Conditional Depletion of TgREMIND abrogates the secretion of GRA and ROP proteins into the PVM. A) In the presence of ATc, GRA1, GRA2, GRA3, GRA4 and GRA5 proteins were all absent from the PVM of the TgREMIND-depleted mutants (right panels). These GRA proteins were present in the PVM of ATc-untreated mutants (left panels, yellow arrows). B) Lack of rhoptry ROP1 protein in the PVM of TgREMIND-depleted mutants (top right and middle panels) while in the untreated TgREMIND mutants, the protein displays typical strong signal staining the PVM (top left and middle panels, yellow arrows). Under the same conditions, the rhoptry ROP7 protein, appears as a punctuated and fragmented signal (bottom right signal, white arrows) compared to the normal compacted signal of untreated mutants (bottom left panel). C) Complementation of iKO-TgREMIND-deficient mutants with ectopic full length TgREMIND-FLAG (Comp-TgREMIND-Flag) restored the normal decoration of ROP1 (middle left panel) and compacted apical signal of ROP7 (middle right panel). The complementation also rescued the PVM staining of GRA1 and GRA5 (bottom left and right panels). Top panels correspond to the co-distribution of TgREMIND-HA and Comp_TgREMIND-FLAG proteins (top left panel), and TgREMIND-HA was depleted from the conditional mutants as expected (top right panel).

**Figure 4.**
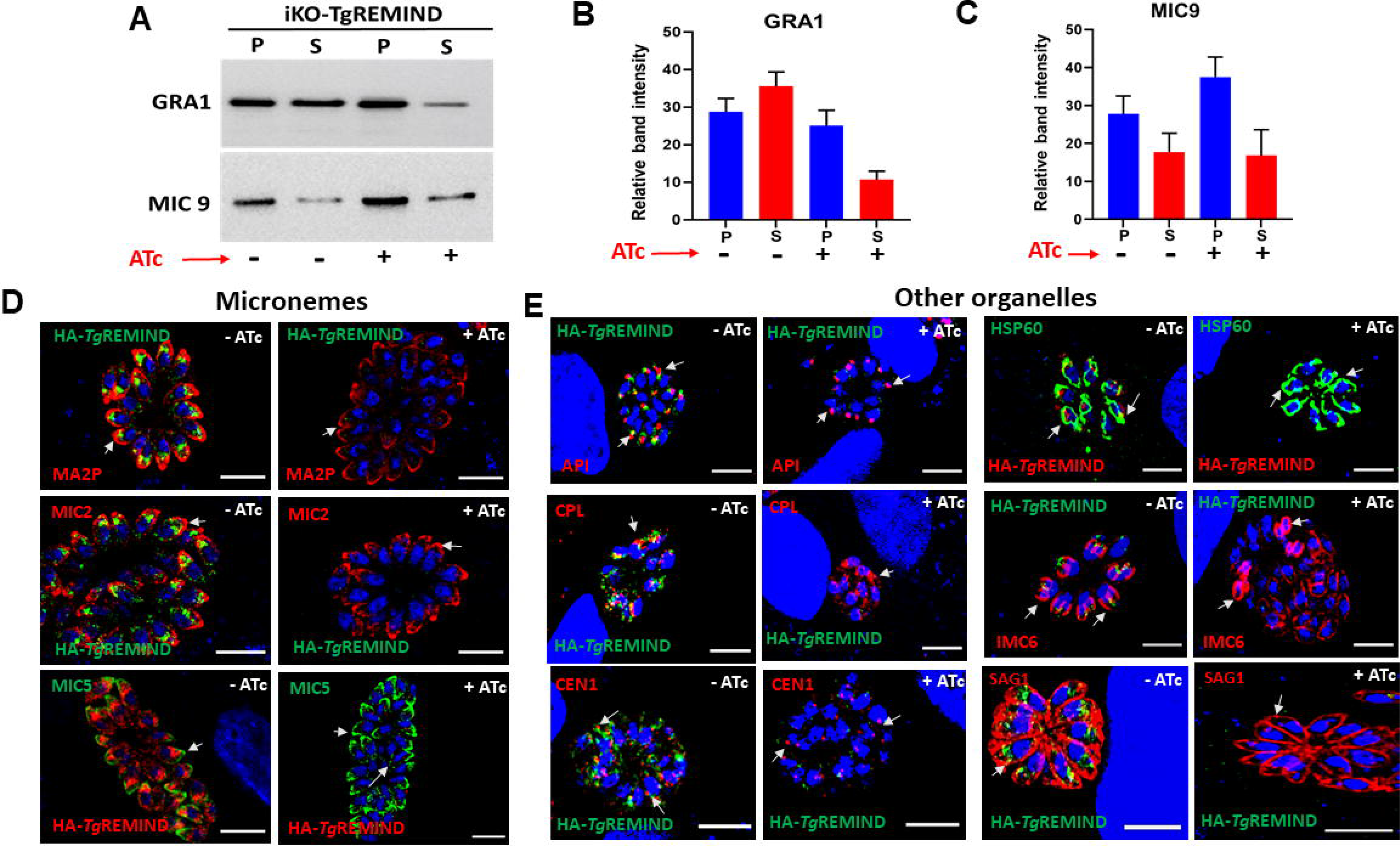
Phenotypic analysis of other organelles in TgREMIND-depleted parasites. A) Illustration of one representative of an immunoblot showing GRA1 (top left panel) and MIC9 (bottom left panel) secretion of untreated *versus* ATc-treated TgREMIND-deficient parasites. B) Quantification of GRA1 and MIC9 secretion from untreated and ATc-treated TgREMIND mutants. Bars, n=3 ± SD. D) Confocal images of TgREMIND depleted parasites show classical and normal pattern of micronemes for anti-MIC (M2AP, MIC2 and MIC5) antibodies, before (left panels) and after (right panels) ATc treatment. E) Subcellular localizations of the apicoplast (Atrx2, Api), mitochondrion (HSP60), cathepsin L-containing compartment (CPL), inner complex membrane (IMC), centriole (CEN1) and plasma membrane (SAG1) markers were unchanged in the TgREMIND-deficient mutant. Bars, 3 µm.

### TgREMIND Is Essential for Organelle biogenesis in *T. gondii*

Multiple vacuoles containing between 16-32 parasites were analyzed by transmission electron microscopy and we found that the vast majority of TgREMIND-depleted mutants were devoid of dense granules (Figures 5C and 5D). In contrast, iKO-TgREMIND parasites contained typical dense granules (Figure 5B, white stars). These data clearly support the absence of GRA proteins secreted in the PVM of intracellular iKO-TgREMIND mutants. In addition, we found that the electron-dense club-shaped rhoptries were present in the TgREMIND-depleted parasites but appeared with a uniformly translucent lumen (Figure 5C, marked Rh) while the untreated iKO-TgREMIND parasites showed a lumen with electron-dense material in both their neck and bulb sub-compartment (Figures 5A and 5B). These ultrastructural data also confirmed the equal presence of the flattened microneme organelles (marked Mn) in the apical end of both iKO-TgREMIND-deficient mutants and TgREMIND-expressing parasites (Figures 5A, 5B, and 5C). We also observed the presence of numerous round and flattened unknown vesicles in the lumen of the PV of TgREMIND-depleted parasites but not in untreated iKO-TgREMIND parasites (Figure 5D, black arrows). However, there were no morphological differences seen for the conoid, Golgi apparatus (marked G), the endoplasmic reticulum (ER), the inner membrane complex (IMC), the plasma membrane, and mitochondrion, as confirmed by the confocal imaging using specific antibodies for these classical organelles or compartments (Figure 4E). Furthermore, complementation of the iKO-TgREMIND mutants rescued the presence of normal electron-dense rhoptries and dense granules that were again detected in these mutants even under the pressure of ATc (Figures 5E and 5F).

**Figure 5.**
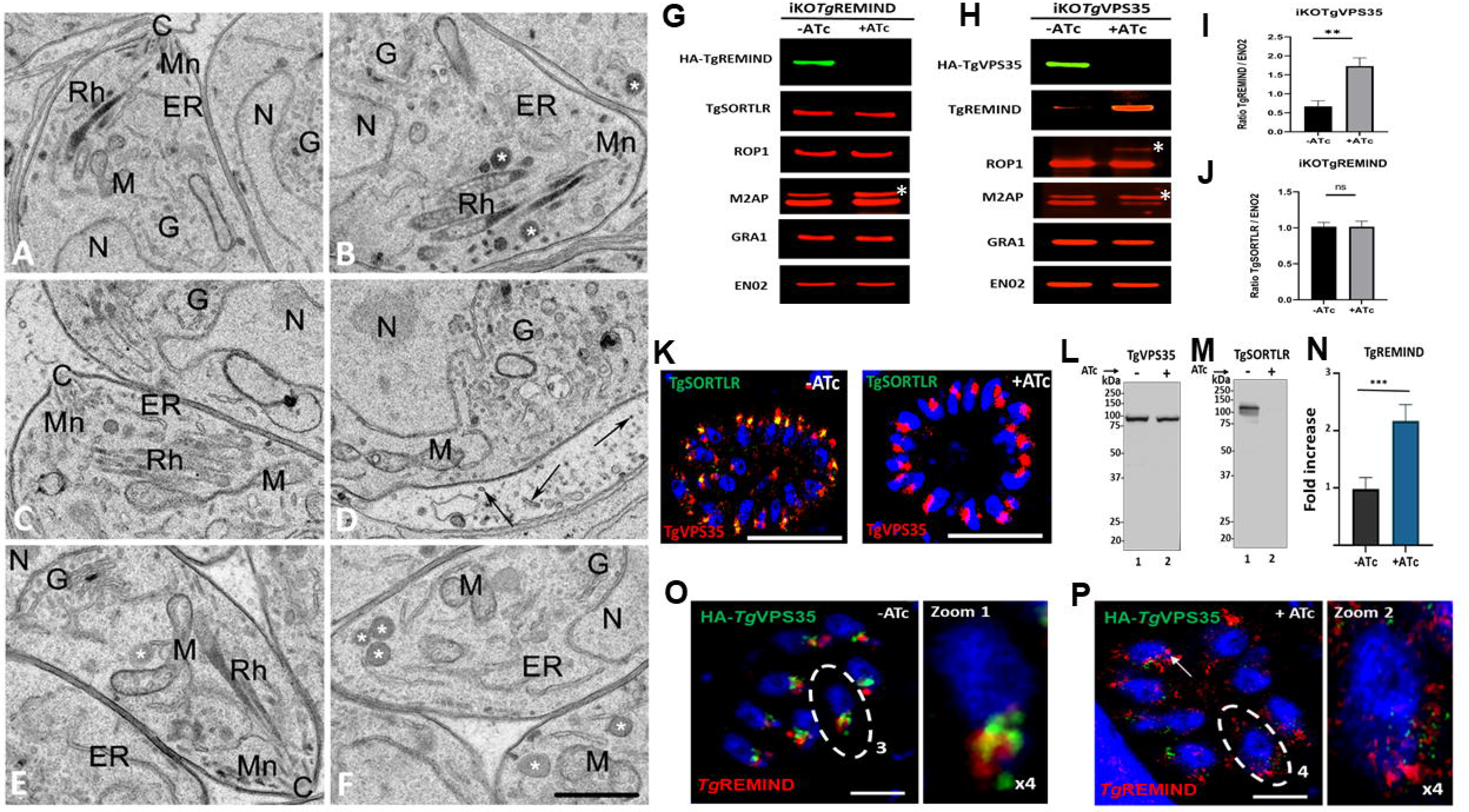
TgREMIND is crucial for biogenesis and functions of dense granules and rhoptries. A and B) Transmission electron micrographs showing typical rhoptries (Rh), dense granules (white stars), micronemes (Mn) in the untreated TgREMIND-deficient mutants. C and D) Transmission electron micrographs of TgREMIND-deficient mutants lacking dense granules and showing translucent rhoptries after 48 h of ATc treatment. E and F) Complementation of TgREMIND-deficient mutants restored the presence of dense granules (white stars) and normal morphology of rhoptries (Rh). C, conoid; ER, endoplasmic reticulum, G, Golgi apparatus, M, mitochondrion and N, nucleus. Bar, 500 nm. G) Immunoblots showing TgSORT, ROP, GRA and MIC protein levels in the ATc-treated and untreated iKO-TgREMIND parasites. ENO2 was used as a loading control. H) Conditional disruption of TgVps35 gene led to an important accumulation of TgREMIND protein level. Inhibition of ROP and MIC processing and maturation was confirmed in TgVps35-deficient parasites. White star corresponds to unprocessed pro-proteins. I) Quantification showing a 3-fold increase of TgREMIND level in TgVps35-deficient parasites. Bars, n=3 ± SD. J) Quantification of TgSORT level was unchanged in TgREMIND-deficient mutants regardless of ATc treatment. Bars, n=3 ± SD. K) Confocal images showing the expression of HA-TgVps35 after gene knock-in into TgSORT-deficient mutants after double homologous recombination in the UPRT locus. IFA were performed using anti-HA and anti-TgSORT antibodies. Bars, 3 µm. L) Immunoblots demonstrating expression of HA-TgVsp35 protein was unchanged after ATc treatment or not, as expected. M) Immunoblots confirming that background of iKO-TgSORT subjected to ATc-treatment led to disappearance of TgSORT protein. N) Quantification of TgREMIND protein level in the knock-in of HA-TgVps35/iKO-TgSORT after co-immunoprecipitation and semi-quantitative mass spectrometry. O) Localization of TgREMIND protein in untreated TgVsp35-deficient mutants using anti-HA and anti-TgREMIND antibodies. Zoom shows that fluorescence of TgREMIND is close to TgVsp35 signal. Bars, 3 µm. P) Localization of TgREMIND in ATc-treated TgVps35-deficient mutants. Bars, 3 µm.

### Cross talk between TgREMIND and the trafficking regulator TgSORT-TgVps35 retromer complex in *T. gondii*

To investigate the role of TgREMIND in ROP and MIC maturation when routed in the ELC, we performed comparative immunoblots using total protein extracts from our conditional iKO-TgREMIND, iKO-TgSORT, and iKO-TgVps35 mutants, respectively. We found no difference in pro-protein processing and maturation of ROP and MIC proteins in TgREMIND-depleted parasites by ATc treatment compared to untreated TgREMIND parasites (Figure 5G, white star). In contrast, we confirmed that ROP and MIC proteins were not processed to maturation in iKO-TgVps35 and iKO-TgSORT mutants (Figures 5H and S5B, see white stars). Still, the level of TgREMIND protein increased more than 3-fold in ATc-treated iKO-TgVps35 parasites compared to untreated mutants (Figures 5H and 5I). No change was seen in TgSORT protein level in iKO-TgREMIND parasites (Figures 5G and 5J). The increased level of TgREMIND protein was also observed in ATc-treated iKO-TgSORT parasites (Figure S5B, see yellow stars). To confirm that TgREMIND protein accumulated at least a 3-fold level under ATc treatment *versus* untreated parasites, we performed co-immunoprecipitation (Co-IP) using cMyc-TgVsp35 as bait (Figure 5N). We checked by immunoblotting that the transgenic TgVsp35-cMyc parasites used for this Co-IP were devoid of TgSORT protein under ATc pressure (Figures 5K and 5M) while the level of TgVps35-cMyc expression was unchanged, as expected (Figure 5L). These data suggest that TgREMIND is a partner of TgVps35, and in the absence of either TgVps35 protein or TgSORT protein, TgREMIND would weakly be used by the parasites to perform organelle biogenesis, thus affecting its turnover. Moreover, in the absence of either TgVps35 protein (Figure 5P) or TgSORT (Figure S5C, right panel), TgREMIND was more dispersed in the cytoplasm of the parasite with the loss of the concentrated signal of the protein on the top of the nucleus where it partially co-localizes with rhoptries and dense granules. The signal of TgVps35-cMyc partially co-localizes with TgREMIND, indicating an amount of TgREMIND protein may be close to cMycTgVps35 (Figure 5O). As expected, TgSORT was also partially co-localized with TgREMIND (Figure S5C, left panels). Spinning Disc Confocal Microscopy coupled to Live SR module revealed that fluorescence signals of TgSORT partially intersected those of TgREMIND, suggesting a cross talk between the TgSORT-TgVps35 retromer complex and TgREMIND at the endosomal-like compartment (Video S1). Interestingly, this spinning disk confocal imaging with a better resolution revealed cytoplasmic vesicles that moved from ER to the TGN/ELC before trafficking to the apical end where rhoptries and micronemes are located (Video S1). Thus, we conclude that functional cooperation between TgREMIND and the TgSORT-TVps35 retromer complex may regulate the formation of vesicles whose contents are essential for the biogenesis of rhoptries and dense granules.

### TgREMIND binds to Inositol Phospholipids and is Required for Secretory Organelle Functions

To gain a deeper understanding of the molecular mechanisms underlying TgREMIND functions in *T. gondii*, we performed computational molecular modeling on the full-length TgREMIND protein (AF-S7W745) using AlphaFold2 (Jumper et al., 2021). The AlphaFold2 predictions indicate that TgREMIND forms a dimer with high confidence (blue color), with the F-BAR domain containing three long kinked α-helices that form a well-packed, crescent-shaped, symmetrical, six-helix bundle. The antiparallel homodimers are indicated as F-BAR (A) and F-BAR (B) (Figure 6A). Additionally, the REMIND domain is also predicted with very high (blue) to high (cyan color) confidence and appears at the two extremities of the antiparallel homodimer (Figure 6A). The secondary structure of the REMIND domain is shown in rainbow-color from the N-terminus to the C-terminus of the domain (Figure 6B). Notably, the REMIND domain is composed of two lobes, the C-terminal lobe starting at strand S7 showing the highest sequence conservation across REMIND family (Figure S3). The lobed structures identified in TgREMIND also exist in human proteins, as predicted by AlphaFold2 with high or very confidence, and they interact with other domains of the proteins (indicated by grey ribbons in Figure S6). In TgREMIND, this region directly contacts the extremities of the F-BAR domains (Figures 6A and 6B). In humans, MADD contains a death domain in the C-lobe (Schievella et al. 1997), which interacts with the Pleckstrin homology (PH)-like domain (Figure S6), known to bind to phosphatidylinositol lipids PI3,4,5P2 and PI4,5P2 (Singh et al. 2021). These findings suggest that the REMIND domain may have a regulatory function on the F-BAR domain. To further demonstrate how the REMIND domain may influence the activity of the F-BAR domain, we complemented the *T. gondii* iKO-TgREMIND mutants by transiently expressing plasmids encoding only the F-BAR or only REMIND domain, and compared this to the full-length TgREMIND protein expression under the same experimental conditions (Figure 6C). When these complemented parasites were incubated with ATc to deplete the endogenous TgREMIND protein, neither the F-BAR domain alone nor the REMIND domain was able to rescue the secretion of ROP1 (Figure 6C, middle and left panels) and GRA1 (Figure 6D, middle, and left panels) into the PVM of the intracellular iKO-TgREMIND-depleted mutants. In contrast, complementation with the full-length FL-TgREMIND perfectly restored ROP1 and GRA1 secretion into the PVM (Figures 6C and 6D, right panels, yellow arrows). These data demonstrate that both F-BAR and REMIND domains are crucial for TgREMIND functions. We then fused the F-BAR, REMIND, and entire TgREMIND to GST at their N-terminus (Figure 6E), which were expressed in *E. coli* and purified (Figures 6F and 6G). Only the full-length TgREMIND was found to strongly bind to PIP2 species, with more intense binding to PI4,5P2, as shown by lipid overlay experiments (Figure 6H). The F-BAR domain alone exhibited extremely weak binding to PIP2 species whereas no binding was observed for the REMIND domain alone (Figure 6H). These results indicate that the presence of the REMIND domain is necessary for the F-BAR domain to efficiently ensure the biological activities of TgREMIND protein.

**Figure 6.**
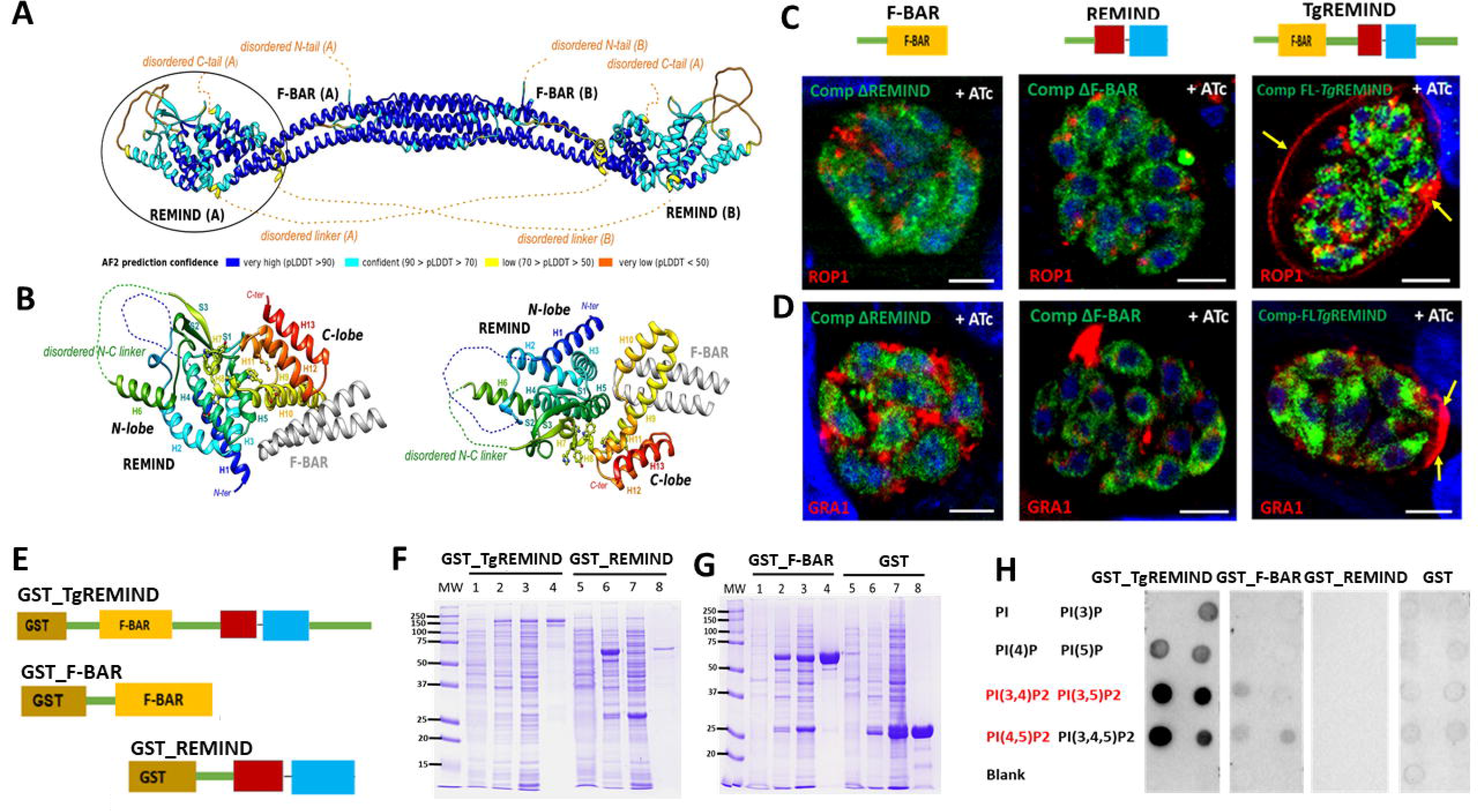
TgREMIND protein binds to PI2P lipids. A) AlphaFold2 model of Tg REMIND showing a dimer, per residue colored according to the pLDDT. The disordered segments are represented with orange, dashed lines. The dimer of F-BAR domains was modeled based on the human formin-binding protein 1 (pdb 2EFL). B) Focus on the REMIND domain (left, encircled domain) and right, with a 90° rotation, colored from blue (N-terminus) to red (C-terminus). C) Schematic representation of FL_TgREMIND-FLAG, F-BAR-FLAG and REMIND-FLAG plasmids for transient transfection and confocal images showing that the N-terminus F-BAR domain alone or the C-terminus REMIND domain cannot restore the presence of ROP1 in the PVM in TgREMIND-deficient mutants (left and middle panels). Only complementation with the full-length TgREMIND protein restored secretion ROP1 in the PVM (right panel). Bars, 3 µm. D) Confocal images show that the N-terminus F-BAR domain alone or the C-terminus REMIND domain cannot restore GRA1 in the PVM of TgREMIND-deficient mutants (left and middle panels). Only complementation with the full-length TgREMIND protein restored GRA1 secretion in the PVM in TgREMIND-deficient parasites under ATc pressure (right panel). Bars, 3 µm. E) Schematic representation of plasmids used to express recombinant GST_TgREMIND, GST_F-BAR and GST_REMIND proteins in *Escherichia coli*. F and G) SDS-PAGE and coomassie blue staining of expression and purification of recombinant of GST_TgREMIND, GST_F-BAR and GST_REMIND. 1, uninduced transformed bacteria; 2, IPTG-induced recombinant bacteria; 3, total extract protein after sonication; 4) GST-affinity purification. H) Lipid overlay experiments using the purified recombinant proteins, and different PIP2 lipid species.

### The Absence of TgREMIND Impairs Parasite Motility and Host Infection

Although transient expression was obtained for all three FLAG epitope-tagged F-BAR, REMIND, and FL-TgREMIND constructs, we could not obtain stable parasite lines containing the plasmid F-BAR domain alone. This confirms that the REMIND domain is required in association with the F-BAR domain to obtain viable and stable complemented parasite lines. Interestingly, transgenic stable parasite lines can be generated using the REMIND domain alone, suggesting that the F-BAR domain alone is toxic to *T. gondii*. Next, we examined the functional role of TgREMIND in host infection by *T. gondii*. In the presence of ATc, iKO-TgREMIND mutants were severely impaired in host cell invasion (Figure 7A), but complementation with FL-TgREMIND restored their invasion capability to that of the untreated mutants (Figure 7A). In contrast, complementation with the REMIND domain alone failed to efficiently complement the iKO mutants (Figure 7A), indicating that both F-BAR and REMIND are also necessary for TgREMIND’s *in vivo* functions. We found no difference between ATc-treated and untreated iKO mutant in intracellular replication (Figure 7B) or in calcium-ionophore induced egress (Figure 7C), indicating that the inability of iKO mutants to invade host cells is not linked to these processes. Whereas the iKO mutants showed a significant number of circular trails, otherwise normal parasite motility in the absence of ATc (Figure 7D, left panel), these mutants were paralyzed in the presence of ATc (Figure 7D, middle panel). As expected, complementation by the full-length FLAG-TgREMIND protein rescued the normal gliding motility despite the presence of ATc (Figure 7D, right panel). Finally, to assess the essentiality of TgREMIND in host infection, we tested the ability of parasites to form plaques on host cell monolayers, which correspond to multiple rounds of parasite entry and egress, ensuring parasite dissemination and host cell lysis. Whereas the wild-type parasites grew normally and developed equal-sized plaques in the absence and presence of ATc, the growth of iKO lines gave rise to normal plaque sizes only in the absence of ATc (Figure 7E). In the presence of ATc, plaque formation was blocked in TgREMIND-deficient parasites (Figure 7E). Complemented iKO-mutants with FL-TgREMIND formed plaques with normal number and size regardless of the presence of ATc (Figure 7E), while complementation with REMIND devoid of the F-BAR domain gave rise to tiny sized plaques, confirming that the absence of F-BAR is detrimental for the *in vivo* functions of TgREMIND in *T. gondii*.

**Figure 7.**
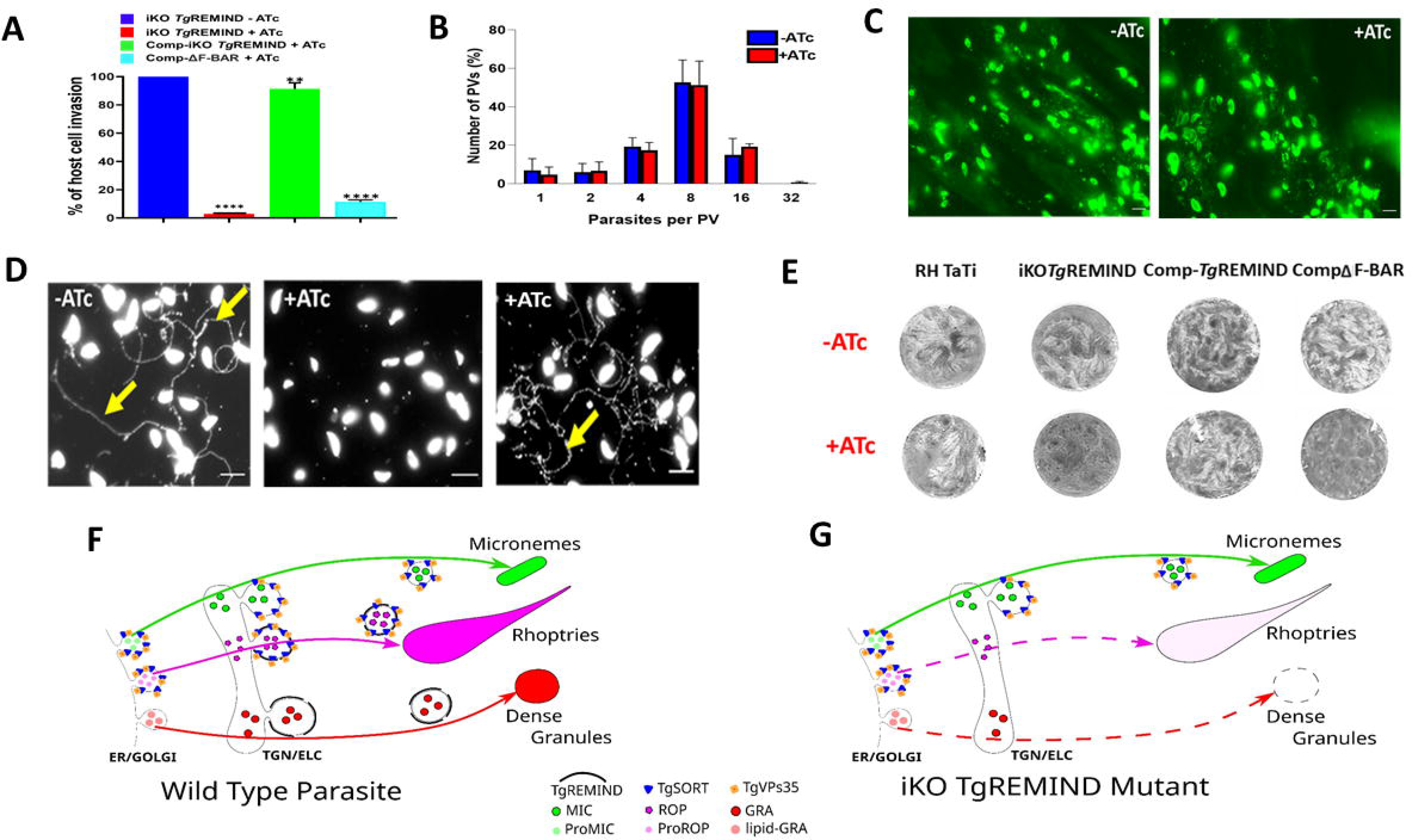
TgREMIND is Essential for Gliding Motility, Host Cell infection and Dissemination. A) Host cell invasion impaired in TgREMIND-deficient mutants after ATc treatment. Complementation with the full-length TgREMIND protein restored host cell invasion whereas complementation with the C-terminus domain alone failed. Bars, n=3 ± SD. B) Identical rate of multiplication of TgREMIND-deficient mutants regardless of ATc treatment. Bars, n=3 ± SD. C) Ca^2+^-ionophore induced egress of TgREMIND-deficient mutants regardless of ATc treatment. D) Gliding (yellow arrows) impaired in the iKO-TgREMIND mutants (middle), rescued complemented mutants (right) and parental parasites (left, yellow arrows). E) Inhibition of plaque formation of TgREMIND depleted mutants (iKO-TgREMIND). Parental (RH TaTi) and complemented (Comp-TgREMIND) parasites shows plaque formation whereas complemented REMIND alone (CompΔF-BAR) parasites shows only tiny plaques. F) Model of TgREMIND functions in biogenesis and secretion of dense granules and rhoptries of *T. gondii*.

## DISCUSSION

In this study, we describe that the TgREMIND protein, which contains both F-BAR and REMIND domains, is indispensable for the proper function of rhoptries and dense granules. TgREMIND belongs to the F-BAR family, and has the dimer crescent or banana shape form that defines this family (Shimada et al., 2007; Henne et al.; 2007). The F-BAR domain in TgREMIND is similar to the FES-CIP4 homology (FCH) domain, which is the archetypal feature of all known F-BAR proteins (Aspenstrom, 1997). These F-BAR-containing proteins are a family of α-helical membrane-binding modules that can detect, induce and regulate membrane curvature, and thereby play essential roles in biological processes, such as endocytosis, exocytosis, regulation of the actin cytoskeleton, cell motility and signaling (Masuda et al., 2006; Shimada et al., 2007). Although the functions of some individual F-BAR proteins have been studied using recombinant proteins and *in vitro* model systems (Simunovic et al., 2015; Snider et al., 2021), but many questions remain unanswered, such as how the accompanying domains located at the C-terminus regulate the precise function of the F-BAR domain and how these proteins are regulated *in vivo*.

Here, we used gene disruption, complementation studies, and predictive structural and biochemical analyses to show that TgREMIND is essential for parasite gliding motility, host infection, and dissemination. In TgREMIND-deficient parasites, rhoptries and dense granules are morphologically and functionally altered. Our results indicate that the presence of the REMIND domain at the C-terminus, in association with the F-BAR domain at the N-terminus, is indispensable for TgREMIND to function properly *in vivo*. This is likely because secretory organelles play crucial roles in all of these phenotypic events. We also showed that TgREMIND is present in membrane-associated components and that only the entire TgREMIND protein strongly interacts with PIP2 lipids *in vitro*. This suggests that the REMIND domain functions together with the F-BAR domain, for binding to PIP2-containing membranes that may play a role in the biogenesis of rhoptries and dense granules. In addition, TgREMIND containing vesicles move from the ER to TGN/ELC before reaching the apical end where rhoptries are located. The presence of actin, myosin, and microtubule proteins suggests that vesicles formed by TgREMIND may traffic along actin filaments or microtubules to reach their destination, similar to F-BAR containing proteins in mammalian cells (Peter et al., 2004; Aspenström, 2009). Notably, the transport and secretion of dense granules depend on actin filaments and myosin F (Heaslip et al., 2016), which were also present in our proteomics data. The requirement of REMIND domain combined with F-BAR to fully complement the TgREMIND deficient mutants and our inability to obtain viable parasite stable lines expressing the F-BAR alone suggest that TgREMIND functions must be tightly regulated in *T. gondii*.

It has been described that DOC2 protein also engages its double Calcium-binding C2 domain with lipids, and membranes to facilitate vesicular trafficking and membrane fusion (Tagoe et al., 2022). *T. gondii* ferlins (FER1 and 2) are responsible for microneme and rhoptry exocytosis, respectively (Farrell et al., 2012). The secretion of rhoptries is also controlled by a set of parasite-specific proteins named rhoptry apical surface proteins (RASP) that cap the extremity of the rhoptry and bind specifically to phosphatic acid (PA) and PI(4,5)P_2_ through a calcium lipid-binding-like domain (C2) and a Pleckstrin Homology-(PH)-like domain (Suarez et al., 2019). TgREMIND, which also binds to PI4,5P2 through its N-terminal F-BAR in association with the C-terminal regulator REMIND domain, is involved in the biogenesis and consequently in secretion of both rhoptries and dense granules. We propose a model in which TgREMIND protein containing F-BAR and REMIND domains forms vesicles that drive anterograde transport containing components such as lipids and proteins destined for rhoptries and dense granules during their biogenesis (Figure 7F). Our data show that deficits in TgREMIND function led to morphological perturbations of dense granules and rhoptries, accompanied by a loss in dense granules and rhoptry exocytosis (Figure 7F). We suggest that TgREMIND is a regulator of membrane bending, curvature, and scission, thus forming vesicles containing components destined for, or recycled from, rhoptries and dense granules. However, the biogenesis and secretion of micronemes depend on Ca2+-dependent control by TgDOC2 or Ferlins (Fareell et al., 2012; Coleman et al., 2018; Tagoe et al., 2022), phosphatidic acid (Bullen et al., 2016) and Rab proteins (Kremer et al., 2013). Taken together, our data support a role for TgREMIND in the biogenesis of only rhoptries and dense granules. The biogenesis of dense granules is less well understood in *T. gondii*, despite advances in electron microscopy. It probably involves several proteins, including TgVps11, TgVps9, and TgVps35 (Morlon-Guyot et al. 2015; Sakura et al. 2016; Sangaré et al. 2016). Our findings demonstrate that TgREMIND, which can bind PIP2 lipids, likely plays a role in membrane curvature and vesiculation, indicating that the molecular basis of dense granule biogenesis in *T. gondii* is more complex than a simple budding of fully matured dense granules from the TGN, as generally reported (Griffith et al., 2022). Our ability to conditionally extinguish TgREMIND expression and complement these mutants will allow functional dissection of regulators of this protein. We plan to further investigate the regulation of TgREMIND, including possible hyper-phosphorylation and/or O-GlcNAcylation of the repeated serine residues at its C-tail. This will help shed more light on the essential sorting mechanism involved in the biogenesis of secretory organelles in *T. gondii*.

## EXPERIMENTAL PROCEDURES

### Growth of host cells and parasite strains

*T. gondii* RH wild-type strain, RHΔku80 (a strain with high homologous integration of transfected DNA; Huynh and Carruthers, 2009) and RHΔku80TaTi parasites, a <u>T</u>rans <u>a</u>ctivator <u>T</u>rap identified inducible anhydrotetracycline (ATc) strain generated in the background of RHΔku80 (Sheiner et al., 2011). These parasites were maintained in monolayer human foreskin fibroblast (HFF) cultured in Dulbecco’s Modified Eagle’s Medium (DMEM) supplemented with 2 mM L-glutamine, 50 µg/mg penicillin/streptomycin and 10% fetal calf serum (FCS) (PAN Biotech, Dutscher, France) in a humid atmosphere under 5% CO_2_ at 37°C as previously described (Sloves et al., 2012).

### Generation of *Tg*REMIND Knockout and Knock-in *T. gondii* Strains

To obtain the conditional iKO-*Tg*REMIND in *T. gondii* RHΔku80-TaTi strain, we used the pyrimethamine-resistant plasmid pG13-D-T7S4 that contains the inducible promoter TeTo7-Sag4. By PCR, we first amplified by PCR, 2 kb of 5’-DNA fragment of *Tg*REMIND, which was cloned into NdeI restriction enzyme site. After verification of the correct orientation using StuI/EcoRV, we cloned the 1.3 kb 3’-DNA fragment of *Tg*REMIND that contains an epitope HA downstream of the inducible promoter. All oligonucleotides used during this work were listed in Table S2 (see Supplemental Information). Fifty µg of this plasmid was linearized with EcoRV and used to transfect 10^7^ tachyzoites of RHΔku80-TaTi that were selected with 2 µM of pyrimethamine before cloning by limiting dilution to obtain the conditional iKO-HA-*Tg*REMIND mutants. We used the pUPRT (Uracil PhosphoRibosyl Transferase) plasmid to complement these mutants, in which either 2 kb of TgREMIND, 1 kb of Tubulin, or 1.5 kb of GRA1 promoter was respectively cloned downstream of the 5’UPRT using AscI/StuI. Then, the full length of *Tg*REMIND (2928bp) was tagged to the epitope FLAG using StuI/AvrII. Finally, the 3’SAG1 untranslated region (350bp) was cloned upstream of the 3’UPRT region using AvrII/PacI. This later plasmid was used to subclone and replace the DNA coding full-length TgREMIND by the 1.15 kb or 1.77 kb of DNA coding, respectively the N-terminus F-BAR and C-terminus REMIND domains with all tagged to the epitope FLAG using StuI/AvrII. DNA sequencing (Genewiz, Germany) using primers listed in Table S2 checked the accuracy of all plasmids. After co-transfection of 10^7^ iKO-HA-*Tg*REMIND parasites of those above linearized, 50 µg of each linearized plasmid to which 10 µg of pSAG1::CAS9-U6::sgUPRT was added to favor a high rate of homologous recombination into UPRT locus (Sidik et al., 2016). Twenty hours later, the co-transfected parasites were incubated with 5µM of 5-fluoro-2’-deoxyuridine (5’ FUDR), and the stable parasite lines obtained were cloned by limiting dilution. After screening by indirect immunofluorescence assays (IFA) using anti-HA or anti-FLAG, the positive clones were selected. The iKO-TgVps35-HA/KI-TgREMIND-cMyc line was generated using the iKO-TgVps35-HA (Sangaré et al., 2016) and pLic-cMyc-TUB5/CAT plasmid (Huynh and Carruthers, 2009) containing 3 kb genomic DNA of TGGT1_259720 gene upstream to stop codon. After linearization of the plasmid by BstZ171, 5×10^6^ parasites were transfected with 50 µg of plasmid, and the stable line was selected with 20 µg of chloramphenicol.

### Immunofluorescence Assays

Paraformadehyde-fixed intracellular parental RHΔku80-TaTi and iKO HA-*Tg*REMIND mutants treated or not with 1.5 µg/ml of anhydrotetracycline (ATc) were used for immunofluorescence assays. The list of monoclonal and polyclonal antibodies used and their dilutions are listed in Table S3 (Supplemental Information). Alexa Fluor secondary antibodies (488 nm, 568 nm, 594 nm) from either Rat, Rabbit, or Mouse were chosen depending on the nature of primary antibodies used at 1:1000 dilutions. In some cases, when a specific primary antibody was not available, we transiently transfected 5×10^6^ tachyzoites with plasmids expressing different markers of subcellular compartments such as ELC (Rab5-HA and Rab7-HA), endoplasmic reticulum (HDEL-GFP), ER/post-Golgi (ERD-GFP), post-Golgi (GalNac-T-GFP) and dynamin-containing compartment (DrpB-YFP(Breinich et al., 2009). The antibodies specific to HA or anti-GFP were used to amplify the fluorescence, then anti-TgREMIND and mouse secondary antibodies were used for co-distribution studies.

### Confocal laser scanning microscopy

The acquisition of fluorescence images were performed using the Leica SP8-X (Leica DMI 6000) microscope with a 63x objective. ImageJ was used for image processing, and Pearson’s coefficient for quantification from 4 to 6 vacuoles, each containing between 4 to 16 parasites. The primary antibodies and dilutions used were listed in Table S3. The secondary antibodies used were either goat anti-mouse, anti-rat or anti-rabbit conjugated to Alexa Fluor 488, 568 or 594 nm (Invitrogen or Thermo Fisher Scientific, France).

### Confocal Spinning Disc Microscopy

IFA images were also obtained using the Inverted Nikon Ti Eclipse Eclipse-E microscope coupled with a spinning disk (Yokogawa, CSU-X1-A1), a 100x plan apo objective (Nikon, NA 1.49, oil immersion), and a sCMOS camera (Photometrics, Prime 95B). An additional “Live-SR” module (GATACA systems) was added to increase two fold the resolution. DAPI fluorescence was detected after a 405 nm excitation (Vortran, 100 mW laser) with a 450/50 bandpass filter (Chroma). Alexa 488 was detected after a 488nm excitation (Vortran, 150mW laser) with a 524/45 bandpass filter (Semrock). Alexa 568 was detected after a 561nm excitation (Coherent, 100mW laser) with a 607/36 bandpass filter (Semrock). The whole system was driven by Metamorph (version 7.7, Molecular Devices).

### Electron microscopy

Intracellular parasites were fixed at room temperature for 1h in 0.1 M sodium cacodylate buffer containing 2.5% glutaraldehyde. Then, samples were kept overnight at 4°C. After washing with the same buffer, samples were fixed again in aqueous 1% osmium tetroxide and 1.5% potassium ferrocyanide. After successive dehydration in ethanol, samples were embedded in Epon (Agar Scientific, AGR1165): 2h in resin mixed with ethanol and 2h with pure resin before polymerization 24h at 60°C. Sections of 70 nm were cut using an ultramicrotome LEICA UC6, collected on grids, and contrasted 15min in 2% aqueous uranyl acetate and 2min in Reynold’s lead citrate.

Ultrastructural images were collected using a JEOL1400 transmission electron microscope at 80kV and camera RIO9.

### Production of Glutathione-*S*-Transferase (GST) Recombinant Proteins and Specific Antibodies

The full length of *Tg*REMIND DNA sequence and those of its N-terminus F-BAR and C-terminus REMIND domains were synthesized and cloned into the pGEX plasmid by GenScript using the restriction enzymes BamHI and NcoI. BL21 DE3 *Escherichia coli* (Biolabs) was used to transform and produce recombinant proteins. First, pre-cultures of 10 ml grown in the presence of ampicillin (100 µg/ml) at 37°C overnight were transferred to one-liter cultures, which were incubated at 37°C for 4 to 5 hours OD_600_ of 0.6. Production of these recombinant proteins was induced with 0.2 mM isopropyl β-D-1-thiogalactopyranoside (IPTG) at 20°C overnight and checked by SDS-PAGE. Next, cultures were centrifuged at 6000xg at 4°C for 15 min. Pellets were frozen at - 80°C for at least 2 h before lysis in 50 mM Tris-HCl pH 8 buffer, 250 mM NaCl, 5% glycerol, and 0.1% Triton X-100 supplemented with 1X protease inhibitors, 1 mg/ml lysozyme, 5 mM DTT and 1 mM Phenylmethylsulfonyl fluoride (PMSF). After incubation for 30 min on ice, bacteria were lysed by 6 cycles of sonication (30 s pulse at 90% and 20 s pause) and centrifuged at 2000 rpm and 4°C. Next, the supernatants were incubated with a GST column overnight at 4°C with 5 mM ATP and 1 mM PMSF. Two series of washes were performed at pH 7.5 with two different washing buffers: 50 mM Tris-HCl, 1 M NaCl, 5 mM DTT, and 50 mM Tris-HCl, 150 mM NaCl, and 5 mM DTT. Finally, recombinant proteins were eluted for 10 min at 4°C using 50 mM Tris-HCl pH 8, 150 mM NaCl, 10 mM GSH (glutathione), 5 mM DTT, and 0.5 mM PMSF buffer. The eluates were dialyzed and concentrated by the Amicon system before SDS-PAGE and Coomassie blue staining. The purified recombinant full-length GST-FL_ TgREMIND, GST_F-BAR and GST_REMIND proteins were used to immunize a group of five Balb/c mice using 50 µg of protein (per mouse) and complete Freund adjuvant. The mice were challenged three times with the same amount of protein prepared in incomplete Freund adjuvant. Following a last boost, the sera were collected and tested by Western blots and IFA. The positive sera specific to each recombinant protein were pooled and purified.

### GST Pull down and Liquid Chromatography-Tandem Mass Spectrometry

To identify *Tg*REMIND partners, 10^9^ tachyzoites of RH strain were lysed for 1 h at 4°C under rotary shaking in a buffer containing 10 mM Tris-HCl, 150 mM NaCl, 1 mM MnCl_2_, 1 mM CaCl_2_, 1% Triton X-100 and protease inhibitors. The lysate was centrifuged at 10000 rpm for 30 minutes at 4°C, and supernatants were incubated with 100 µg of each recombinant protein GST-*Tg*REMIND, GST-F-BAR, GST-REMIND coupled to GST beads or with GST alone (negative control) at 4°C overnight. Five washings were performed with lysis buffer before elution, and SDS-PAGE analyses. Protein bands were excised and processed for LC-MS/MS.

### Cell Fractionation

Extracellular parasites that were freshly egressed of wild type *T. gondii* RHΔku80 (10^8^ tachyzoites) were purified to remove HFF debris, pelleted, and washed once with PBS as described (Breinich et al., 2009). Each pellet was resuspended with 300 µl of each different lysis buffer as follows: PBS 1x, 2% Triton X-100 in 1x PBS, 1M NaCl in 1x PBS, and 0.1 M Na_2_CO_3_ in PBS 1x at pH 11.5. After lysis by three series of rapid freezing in liquid nitrogen and thawing in a water bath at 37°C, homogenates were obtained by sonication. The different homogenates underwent ultracentrifugation at 100,000xg at 4°C to separate into different soluble and insoluble fractions. Fractions equivalent to 1×10^7^ parasites were analyzed by SDS-PAGE followed by immunoblots using specific anti-TgREMIND, anti-Glycosyl PhosphoribosyI Inositol (GPI) anchored SAG3 and anti-G6-PI (glucose 6-phosphate isomerase) antibodies.

### Microneme Secretion and Dense Granule Excretion

Freshly egressed (2×10^8^ parasites) ATc-treated (1.5 µg/ml) or untreated iKO-TgREMIND mutants for 48 h were harvested by centrifugation at 600 x g for 10 min at room temperature. The samples were washed twice with prewarmed at 37°C of intracellular buffer containing 5 mM NaCl, 142 mM KCl, 1 mM MgCl_2_, 2 mM EGTA, 5.6 mM glucose and 25 mM HEPES, pH 7.2). After centrifugation, parasites were resuspended in DMEM supplemented with 2 mM glutamine containing 500 µM of propranolol or not and incubated for 20 min at 37 °C (Bullen et al., 2016). Parasites were centrifuged at 1000 × g, 4 °C for 5 min. Pellets were washed once in PBS and resuspended in Laemmli buffer. Supernatants were centrifuged at 2000 × *g* for 5 min, 4 °C, and supernatants used as ESA (Excreted/Secreted Antigen) were also suspended in 2XLaemmli buffer. Pellets and ESA samples were run on SDS-PAGE and analyzed for the presence of dense granule GRA1 and microneme MIC9 by Western blots.

### Co-immunoprecipitations, SDS-PAGE and Western Blots

For co-immunoprecipitations, 2×10^8^ parasites from transgenic knocked in of cMyc-TgVps35 into iKO-TgSORT mutants freshly egressed after ATc-treatment or not for 48h were purified and extracted with Triton X-100 as described (Fauquenoy et al., 2011). Binding protein complexes of co-immunoprecipitation or total protein extracts of purified parasites were electrophoresed on SDS-PAGE, transferred at 80 V for 1 h 15 min onto a nitrocellulose membrane in a cold buffer containing 25 mM Tris-HCl, 190 mM glycine and 20% methanol in a cold room. After 5 min of 0.2% Ponceau red staining, water-rinsed membranes were saturated with 5% skim milk prepared in TNT buffer (15 mM Tris-HCl pH 8; 140 mM NaCl; 0.05% Tween-20) for 30 min at room temperature. The primary antibodies used are listed in Table S3. The secondary peroxidase-coupled antibodies (Sigma, France) or Alexa fluor 647 nm (Invitrogen, France) or Plus 800 nm (Thermo Fisher Scientific) antibodies (Invitrogen, France) prepared in TNT were incubated for 1 h at room temperature. Following three washes, the blots were revealed with ECL (Enhanced ChemiLuminescence Plus Western Blotting Detection System, Amersham Biosciences) whereas the Alexa fluor antibodies were directly scanned using VILBER Fusion FX imaging apparatus (France).

### Gliding Assay and Host Cell Invasion

Trail deposition assays were performed as previously described (Fauquenoy et al., 2008). ATc-treated iKO-TgREMIND and ATc-treated complemented Comp-KO-TgREMIND parasites for 48h were purified, and pellets were resuspended with incomplete DMEM medium supplemented with 10 mM HEPES and 1 mM EGTA. Parasites (5.10^6^ tachyzoites) were plated on glass coverslips previously treated with 50% FCS at 37°C overnight. After 20 minutes of incubation at 37°C under 5% CO_2_, parasites were fixed with 4% PFA. Gliding trails were observed under a Zeiss fluorescence microscope after staining with SAG1 antibodies. For host cell invasion, iKO-TgREMIND, Comp-full length TgREMIND_FLAG and Comp-REMIND_FLAG were treated or not with ATc for 48h. Freshly egressed parasites were used to infect confluent HFF cells and processed as described (Sangaré et al., 2016).

### Plaque assays

Plaque assays were carried out as previously described (Sloves et al., 2012), except that the coverslip containing confluent HFF cells were infected with 500 iKO-*Tg*REMIND, Comp-iKO*Tg*REMIND, Comp-REMIND, and RHTaTi. After 3h post-infection, 1.5 µg/ml of ATc was added or not to these infected HFF coverslips and incubated for seven days. Infected HFF cells were fixed with cold methanol on ice for 10 min before staining with crystal violet at room temperature for 10 min. After three washes with distilled water, the coverslips were air-dried before imaging.

### Bioinformatics analyses

Sequences were downloaded from the UniProt database (UniProt Consortium, 2021). AlphaFold2 (AF2) (Jumper et al. 2021) models were extracted from the AFDB database (Varadi et al. 2022). Sequence similarities were initially searched against UniProt using PSI-BLAST (Altschul et al. 1997). The jackhammer software (Finn et al. 2015) was also used to query the initial REMIND domain alignment and target the Uniprot reference proteomes with default parameters. HH-PRED (Zimmermann et al. 2018) was used to search protein domain and structure databases. A comparison of 3D structures against those included in the Protein Data Bank (PDB) was performed using DALI (Holm 2022). 3D structures were visualized with the UCSF Chimera package (Pettersen et al. 2004). Sequence alignment was rendered using ESPript (Robert and Gouet 2014).

### Statistical analysis

We analyzed all the data with GraphPad9 software, and statistical studies were performed using ANOVA one-way or two-way followed by Tukey’s multiple comparisons tests.

### Additional Information and accession number

The mass spectrometry proteomics data generated by nanoLC-MS/MS analyses have been deposited to the ProteomeXchange Consortium via PRIDE partner repository with the dataset identifier and accession code: PXD025049 and 10.6019/PXD025049.

## ACKNOWLEDMENTS

We thank our colleagues who generously provided invaluable antibodies for this study. In addition, we are grateful to Omar Ndao and Scott Thomas for technical assistance, members of the Gif Imaging Facility (I2BC, CNRS): Cynthias Dupas for electron microscopy, Romain Le Bars and Sandrine Lecart for the training and help on confocal microscopy; and other members of the Lab: Ariane Honfozo and Mahendra Jamdhade for their fruitful discussions. This work was supported by a Ph.D. fellowship from the Campus France, I2BC (CNRS) and WAEMU to R.H, and by financial support from the Agence Nationale de la Recherche grant N°ANR-19-CE44-0006 to ST.

## AUTHOR CONTRIBUTIONS

The author(s) have made the following declarations about their contributions: Performed the experiments: RH, LOS, TDA, AD, CB

Contributed reagents/materials/analysis tools: YH, CMA, LAF, CS, TBF, IC, JPJS Performed data analysis: LOS, TDA, CB, CS, ST

Conceived and designed the experiments: ST

Wrote the paper: RH, LOS, YT, IC, JPJS, ST

Supervising the work: ST

All authors approve the content and submission of the paper.

