## Supplementary figures and images for "*Toxoplasma* Membrane Inositol Phospholipid Binding Protein TgREMIND Is Essential for Secretory Organelle Function and Host Infection"

### 1.JPG

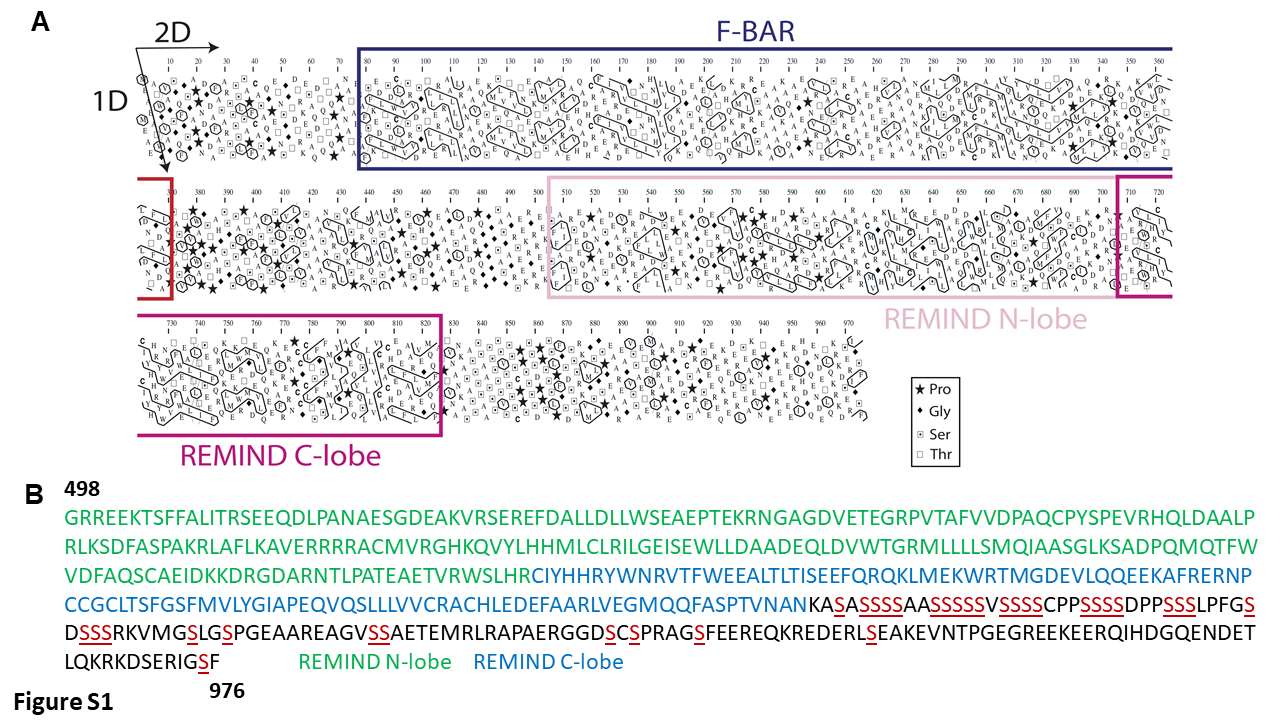

### 2.JPG

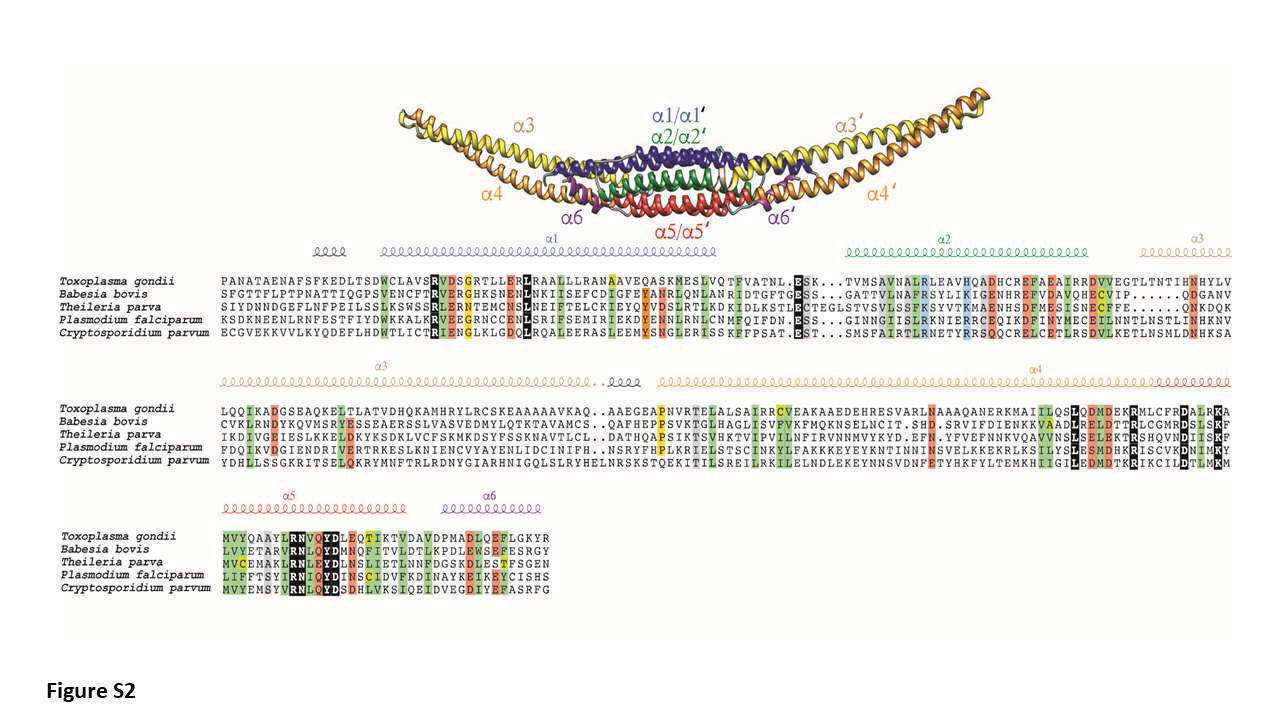

### 3.JPG

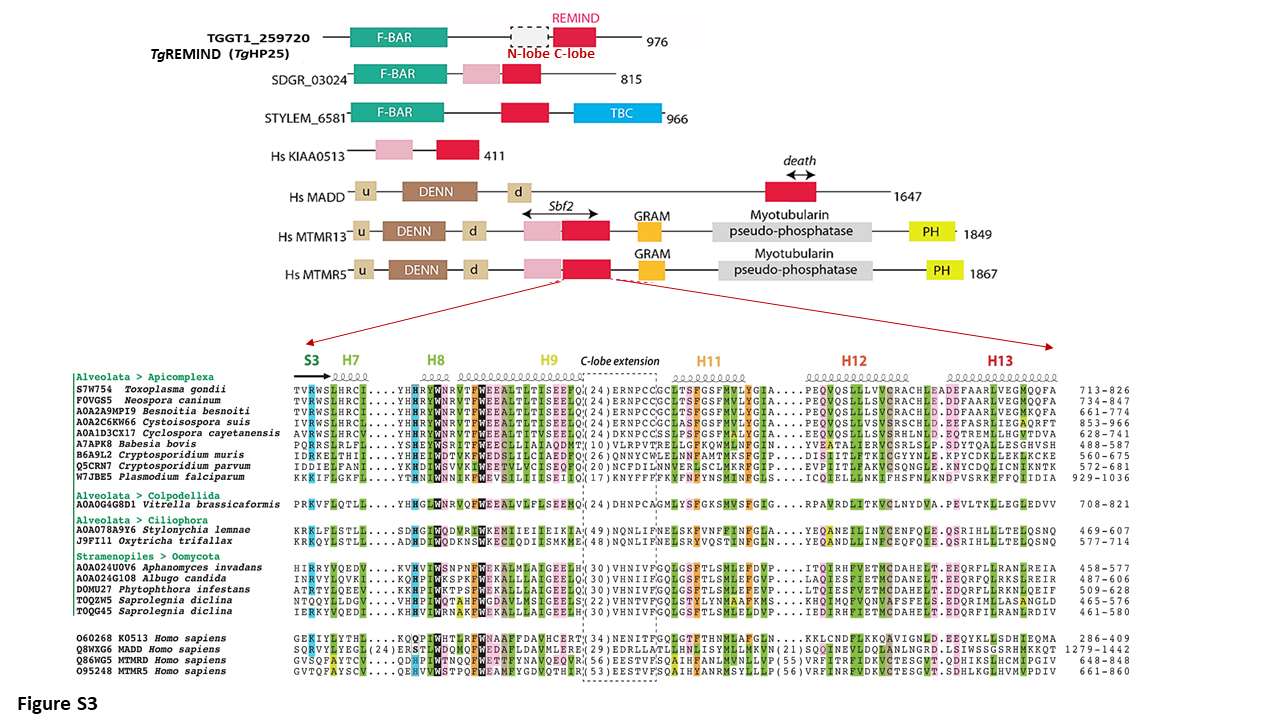

### 4.JPG

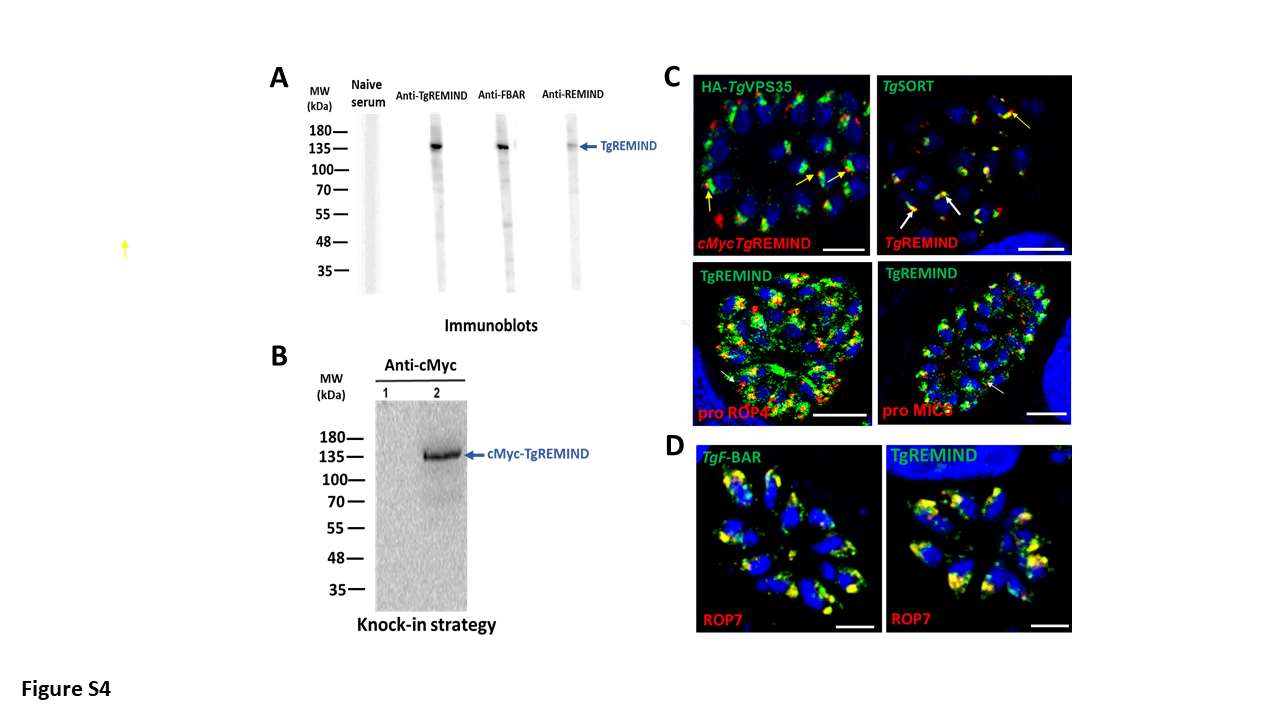

### 5.JPG

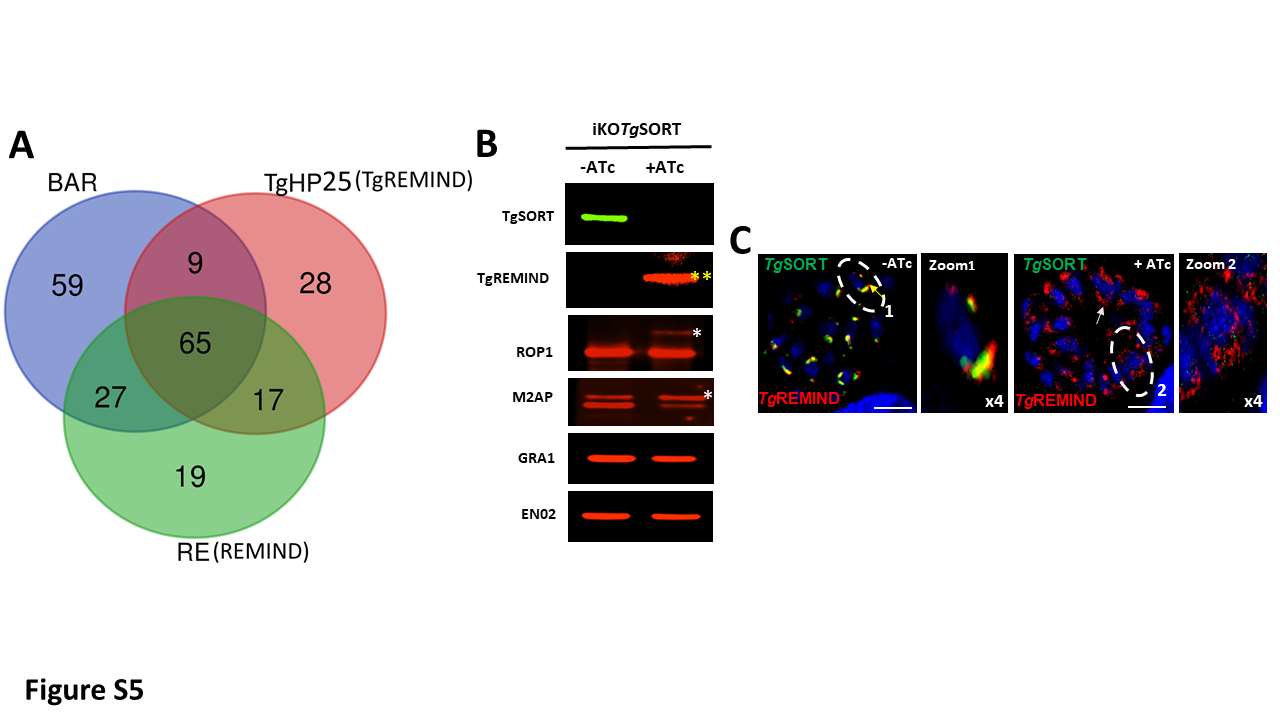

### 6.JPG

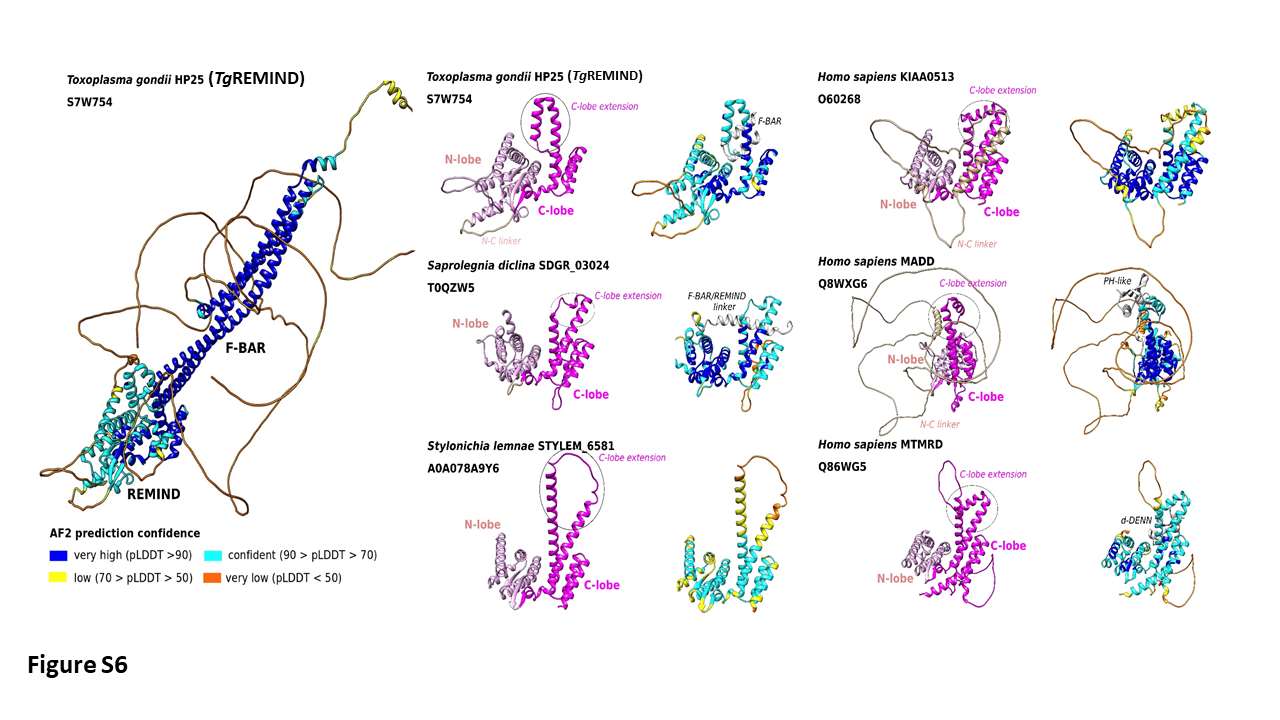

### Page1_Thumbnail.JPG

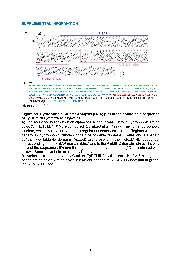

### Video S1

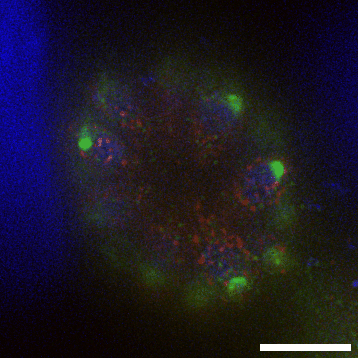
