## Supplementary material for "*Toxoplasma* Membrane Inositol Phospholipid Binding Protein TgREMIND Is Essential for Secretory Organelle Function and Host Infection": Table S2

**Table S2**: Oligonucleotides used for PCR and restriction enzymes for cloning

| **Names** | **5' Sequences 3'** | **Restriction sites** | **Purpose** |
|---|---|---|---|
| iKO_*Tg*REMIND_5'_F | CCGGCATATGACAGCTATTCCGGGCTGAGGTGAGCTCA | NdeI | iKO-*Tg*REMIND |
| iKO_*Tg*REMIND_5'_R | CCGGCATATGTCGTGAGAACGGCTTCAGCACGGAGTTCTT | NdeI | iKO-*Tg*REMIND |
| iKO_*Tg*REMIND_3'_F | CCGGAGATCTATGTACCCATACGATGTTCCAGATTACGCTGAAGCGGAGGCCACC TG | BglII | iKO-*Tg*REMIND |
| iKO_*Tg*REMIND_3'_R | CCGGCCTAGGAGCGTTGACGGCAGACATGAC | AvrII | iKO-*Tg*REMIND |
| Locus_*Tg*REMIND_F | ACACGCGTGTTTGACAGTCTCACAC | | Integration of pDTS4-*Tg*REMIND |
| DHFR_Int_R | GGCGTTGAATCTCTTGCCGACTGATGGAGAGGGAAGTCC | | Integration of pDTS4-*Tg*REMIND |
| Compl_FL*Tg*REMIND_F | TATTAGGCCTATGGAAGCGGAGGCCACCTGGGGGGTGGCGGGGTCC | StuI | Complementation for FL and *Tg*F-BAR |
| Compl_FL*Tg*REMIND_R | CCCTTAATTAACCCCTAGGTTATCACTTATCGTCATCGTCTTTGTAATCAAAAGAG CC | AvrII | Complementation for FL and *Tg*SBF2 |
| Compl_*Tg*F-BAR_R | GCATCCTAGGTCACTTATCGTCATCGTCTTTGTAATCTGGCGTCGACCCCGGCGC | AvrII | Complementation |
| Compl_*Tg*REMIND_F | GCATAGGCCTGCCCACGGCTCTTCGGGGTC | StuI | Complementation |
| Compl_*Tg*GRA1prom_F | CATTGGCGCGCCCTCACCTTTGGCTCAATCCTA | ASCI | Complementation |
| Compl_*Tg*GRA1prom_R | CGACAGGCCTCTTGCTTGATTTCTTCAAAGAACAA | StuI | Complementation |
| Compl_*Tg*SAG1_3'_F | TATTCCTAGGTCACCGTTGTGCTCACTTCTCAAATC | AvrII | Complementation |
| Compl_*Tg*SAG1_3'_R | CCCTTAATTAA*CCC*GACGATCGCCCATCGGGGTCGTGA | PacI | Complementation |
| KI_*Tg*REMIND-cMyc_F | TACTTCCAATCCAATTTAATGCTTTTTCGCGCTGATTACGAGAAGCGAGG | | Knock-in *Tg*REMIND |
| KI_*Tg*REMIND-cMyc_R | TCCTCCACTTCCAATTTTAGCAAAAGAGCCGATCCTCTCCGAGTCTTT | | Knock-in *Tg*REMIND |

| | | | |
|---|---|---|---|
| TgREMIND Sense primer 1 | CTCGCTGCGTCGTTCTGAGTC | | DNA sequencing |
| TgREMIND Sense primer 2 | GGACGAGAAGCGCATGC | | DNA sequencing |
| TgREMIND Sense primer 3 | CCGAAGGCAGACCCGTCA | | DNA sequencing |
| TgREMIND Sense primer 4 | TGGGGAGTCTTGGGTCTCC | | DNA sequencing |
| TgREMIND Antisense primer 1 | ACCGGTTCGACTAGAACAACT | | DNA sequencing |
| TgREMIND Antisense primer 2 | GAGGCGGGCTGCGAATTC | | DNA sequencing |
| TgREMIND Antisense primer 3 | TCTCAGATCGCACCTTCG | | DNA sequencing |
| TgREMIND Antisense primer 4 | CAGCGCCGAATCGCCGAC | | DNA sequencing |
