## Supplementary material for "*Toxoplasma* Membrane Inositol Phospholipid Binding Protein TgREMIND Is Essential for Secretory Organelle Function and Host Infection": Table S3

**Table S3**: List of monoclonal and polyclonal antibodies used

| Name | Species | Dilution for IFA | Dilution for Western Blot | Origin |
|---|---|---|---|---|
| Anti *Tg*REMIND | Mouse | 1:500 | 1/500 | Tomavo Lab (this study) |
| Anti F-BAR | Mouse | 1:500 | 1:500 | Tomavo Lab (this study) |
| Anti REMIND | Mouse | 1:500 | 1:500 | Tomavo Lab (this study) |
| Anti *Tg*SORT | Rat | 1:500 | 1:1000 | Tomavo Lab |
| Anti HA | Rat | 1:250 | 1:1000 | Cell Signaling |
| Anti HA | Rabbit | 1:250 | 1:1000 | Cell Signaling |
| Anti Flag | Rabbit | 1:1000 | 1:1000 | Cell Signaling |
| Anti cMyc | Mouse | 1:500 | 1:500 | Thermo Fisher |
| Anti ROP1 | Mouse/Rabbit | 1:500 | 1:1000 | Dubremetz Lab |
| Anti ROP4 | Mouse | 1:500 | NT | Dubremetz Lab |
| Pro ROP4 | Rabbit | 1:500 | NT | Gary Lab |
| Anti ROP5 | Rabbit | 1:1000 | 1:1000 | Sibley Lab |
| Anti ROP7 | Mouse | 1:500 | 1: 1000 | Dubremetz Lab |
| Anti GRA1 | Mouse | 1:500 | 1:1000 | BIOTEM, France |
| Anti GRA2 | Mouse | 1:500 | NT | BIOTEM, France |
| Anti GRA3 | Rabbit | 1:500 | NT | Dubremetz Lab |
| Anti GRA4 | Mouse | 1:500 | NT | Dubremetz Lab |
| Anti GRA5 | Mouse | 1:500 | NT | BIOTEM, France |
| Anti MIC2 | Mouse/Rabbit | 1:500 | 1:1000 | Carruthers Lab |
| Anti MIC5 | Rat/Rabbit | 1:500 | 1:1000 | Carruthers Lab |
| Anti M2AP | Rabbit/Rat | 1:500 | 1:1000 | Carruthers Lab |
| Anti proMIC5 | Rat | 1:500 | NT | Carruthers Lab |
| Anti MIC9 | Rabbit | 1:500 | NT | Soldati Favre Lab |
| Anti VP1 | Rabbit | 1:500 | NT | Moreno Lab |
| Anti CPL | Rabbit | 1:500 | NT | Moreno Lab |
| Anti HSP60 | Rabbit | 1:500 | NT | Tomavo Lab |
| Anti API (Trx) | Mouse | 1:500 | NT | Parsons Lab |
| CEN1 | Rabbit | 1:500 | NT | Gubbels Lab |
| IMC6 | Rabbit | 1:500 | NT | Bradley Lab |
| Anti SAG1 | Mouse | 1:500 | NT | Dubremetz Lab |

**NT= not tested**
